# Mapping the transcriptional landscape of algal resistance to viral infection reveals a core expression program

**DOI:** 10.1101/2025.04.29.650402

**Authors:** Talia Shaler, Amir Fromm, Daniella Schatz, Shifra Ben-Dor, Ester Feldmesser, Assaf Vardi

## Abstract

- Algal blooms and their demise by viruses drive global-scale ecological processes in the ocean. These blooms form the foundation of marine food webs, regulate microbial communities, and shape biogeochemical cycles. Although algal populations are constantly infected by viruses, resistant subpopulations frequently emerge after the infection. Yet, antiviral molecular mechanisms of marine microalgae are underexplored.
- We used a model system of the ubiquitous microalga *Emiliania huxleyi* and its giant virus, *E. huxleyi* virus (EhV), to study how resistant populations evolve and to identify their transcriptional programs. We generated a detailed temporal transcriptomic dataset during a viral infection, covering the stages of lysis and the recovery of a resistant subpopulation.
- Viral infection triggered prominent transcriptome changes to support viral propagation, followed by a unique transcriptional response in resistant cells. Both infected and resistant cells highly expressed innate immune response genes, notably those with Toll/interleukin-1 receptor (TIR) domain. Additionally, resistant cells expressed genes involved in membrane-bound glycan remodeling, sphingolipid metabolism, and nutrient uptake.
- Using comparative transcriptomics across diverse resistant *E. huxleyi* strains, we identified a core group of resistance-related genes, providing a set of gene markers to detect this rare phenotype during the host-virus arms race in algal blooms.

## Introduction

Phytoplankton, unicellular photosynthetic microorganisms, are key primary producers that form extensive oceanic blooms (Winter *et al*., 2006; Sarkar, 2018; Schleyer & Vardi, 2020). The blooms are often terminated by lytic virus infections (Bratbak *et al*., 1993; Tarutani *et al*., 2000; Martínez Martínez *et al*., 2007; Vardi *et al*., 2012; Laber *et al*., 2018), and the algal biomass is released in a process known as the viral shunt (Wilhelm & Suttle, 1999). This flux of nutrients and metabolites is a key ecological process that fuels marine food webs, microbial communities (Buchan *et al*., 2014), biogeochemical cycles (Simó, 2001; Rost & Riebesell, 2004; Taylor *et al*., 2017), and carbon export to the deep ocean. Although algal blooms demise, they recur annually, and resistant cells were isolated during the demise phase of the blooms (Tarutani *et al*., 2000; Schleyer *et al*., 2023), indicating a stable coexistence of hosts and viruses in the environment (Waterbury & Valois, 1993; Carlson *et al*., 2022). Laboratory experiments in several microalgae species show that resistant populations emerge after viral infection (Thyrhaug *et al*., 2003; Tomaru *et al*., 2009; Thomas *et al*., 2011; Frada *et al*., 2017; Joffe *et al*., 2024). Therefore, studying how resistance to viral infection is acquired in natural populations is key to understanding bloom dynamics and their ecological consequences.

In the host-virus evolutionary arms race, host cells develop multiple layers of defense, and viruses evolve counter-defenses to overcome them. Defense mechanisms can act at the cell surface level to prevent viral attachment and entry by mutating membrane receptors (Stoddard *et al*., 2007; Avrani *et al*., 2011; Kolan *et al*., 2024). Alternatively, they can act intracellularly by interfering with various stages of the viral life cycle, such as gene transcription, translation, genome replication, or virion packaging. In response, viruses evolve to overcome host defense in many ways, from mutation-based evasion (Stokar-Avihail *et al*., 2023) to direct inhibition of immune proteins (Leavitt *et al*., 2022; Yirmiya *et al*., 2025) and restoration of molecules depleted by defense systems (Osterman *et al*., 2024).

New antiviral defense systems are constantly being discovered in bacterial model systems like *E. coli* (Doron *et al*., 2018; Millman *et al*., 2022), with some even functionally conserved in human immune systems (Wein & Sorek, 2022; Bernheim *et al*., 2024). However, for marine microalgae, where host-virus dynamics have broad-scale ecological implications but genetic tools and accurate gene annotations are not always on hand, most defense strategies remain underexplored. Using homology to known defense systems is a common practice for detecting resistance mechanisms, but it may not fully capture the diversity of algal resistance. For example, a recent study tested two putative defense systems encoded by a resistant strain of the cyanobacterium *Synechococcus*. Surprisingly, they did not protect against a cyanophage infection (Zborowsky *et al*., 2025). Instead, resistance was conferred by a constitutive absence of specific tRNAs in the host cells, preventing the translation of key cyanophage proteins. Therefore, elucidating the molecular mechanisms contributing to algal resistance phenotypes requires comparative approaches and functional assays.

To explore genes involved in algal resistance to viral infection, we used the bloom-forming *Emiliania huxleyi* and its specific virus, *E. huxleyi* virus (EhV), a lytic nucleocytoplasmic large DNA virus (NCLDV). When EhV infects *E. huxleyi* cells, it reprograms the cell transcriptome and metabolome into a virocell state that functions to support viral replication (Forterre, 2013; Rosenwasser *et al*., 2014; Ku *et al*., 2020). Infection ultimately leads to cell lysis and bloom demise (Bratbak *et al*., 1993; Wilson *et al*., 2002; Martínez Martínez *et al*., 2007). But, under laboratory conditions, infection is followed by cell regrowth and the recovery of a resistant population (Thyrhaug *et al*., 2003; Joffe *et al*., 2024). This resistant population exhibits a spectrum of resistance levels, enabling a dynamic balance that supports host-virus coexistence (Joffe *et al*., 2024).

Resistance in *E. huxleyi* was previously associated with morphological changes, increased ploidy levels (Frada *et al*., 2017), and different glycosphingolipids (GSL) compositions (Hunter *et al*., 2015; Schleyer *et al*., 2023). Comparative transcriptomics of resistant and sensitive *E. huxleyi* strains identified more than 18,000 differentially expressed (DE) genes (Feldmesser *et al*., 2021). The high number of DE genes makes it difficult to understand which genes contribute directly to the resistance phenotype and which are linked to any existing strain variability, such as morphology or cell physiology. In the current work, we used dual transcriptomics of *E. huxleyi* and EhV in high temporal resolution to explore the molecular basis of the transition from sensitivity to resistance of the algal host to viral infection. Our findings outlined a unique transcriptomic profile of resistant cells observed across multiple resistant *E. huxleyi* strains. This core group of resistance-related genes includes those involved in remodeling membrane-bound glycans, sphingolipid metabolism, and nutrient uptake. Additionally, it involves genes with domains related to innate immune responses, such as Toll/interleukin-1 receptor (TIR) domains. These core resistance-related genes offer a set of potential markers for identifying *E. huxleyi* resistance in complex natural populations.

## Results and Discussion

### Host cells recovered from viral infection and acquired resistance

To investigate the host cellular response that facilitates resistance to viral infection, we infected *E. huxleyi* (strain RCC6946) with *E. huxleyi* virus (EhV; strain EhVM1) and sampled the cultures over a detailed time course across various phases of interactions. This host-virus pair was isolated from induced blooms of *E. huxleyi* in the Fjords of Norway and presents an ecologically relevant host-virus model system (Fig. 1a).

**Figure 1.**
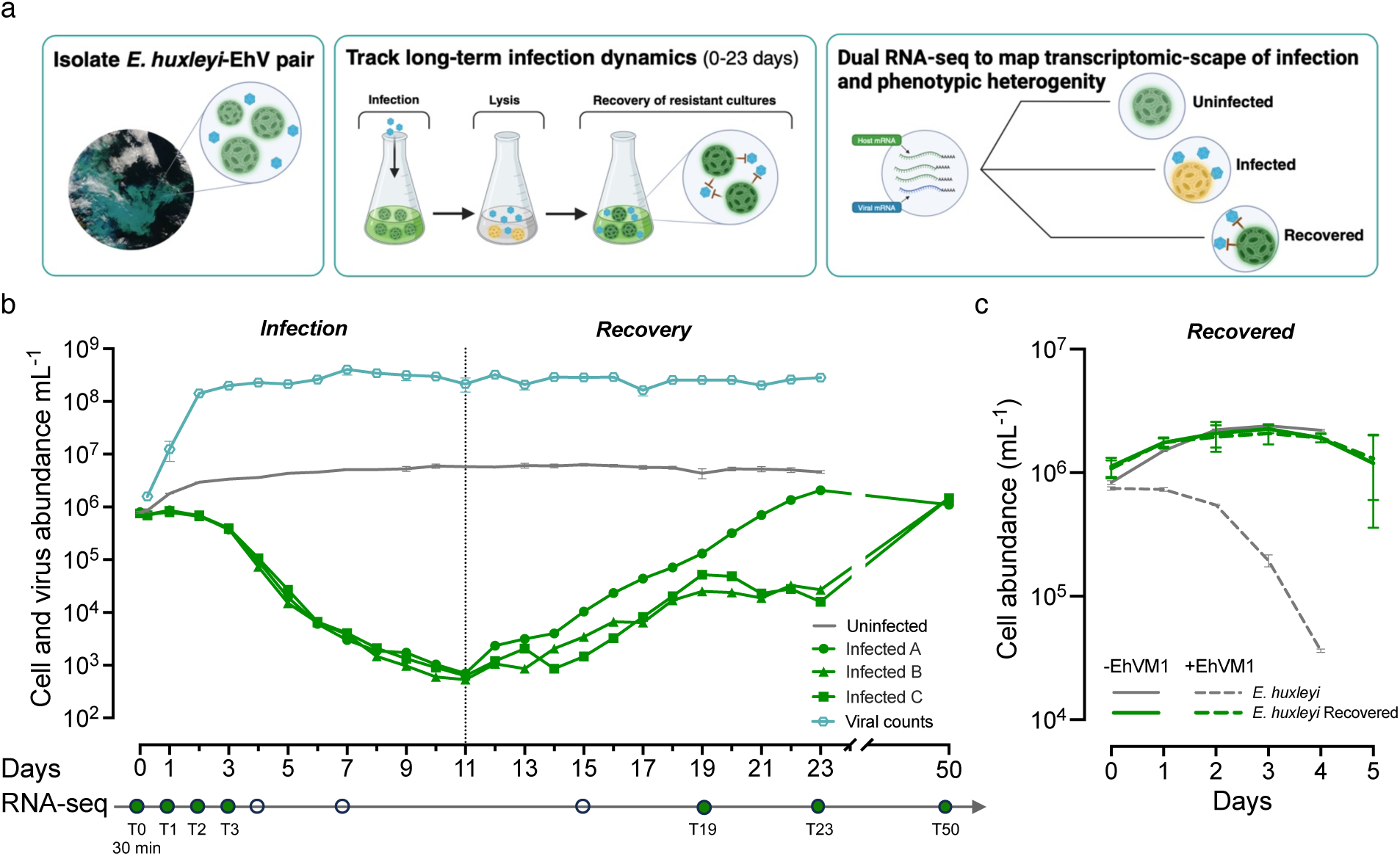
Infection dynamics of *E. huxleyi* and EhV resulted in the recovery of resistant populations. (a) Schematic representation of the experimental system. *E. huxleyi* RCC6946 and EhVM1 strains were isolated from the natural environment. Upon lab-controlled viral infections, *E. huxleyi* populations lyse, but a small number of cells survive and form a new population in the presence of lytic viruses. During the infection process and recovery, the population was sampled for dual RNA-seq, which captures cellular and viral gene expression to characterize the different phenotypes of the population transcriptionally. (b) Infection dynamics of *E. huxleyi* with EhVM1 (MOI = 0.075:1) resulted in the recovery of resistant populations in the presence of infective viruses. Cell counts of uninfected replicates (grey) and viruses in infected cultures (light blue) are represented as mean ± SD (n = 3). For the infected samples (green lines), each replicate is detailed. Error bars may be smaller than the symbol size. The bar below outlines which time points were sampled for RNA-seq (filled circles); empty circles represent time points where live cell concentration was too low to produce high-quality RNA libraries. (c) Growth of the recovered *E. huxleyi* population was not affected by re-inoculation with EhV. Schematic icons were created with BioRender.com.

*E. huxleyi* exponential cultures were infected with EhVM1 and monitored by flow cytometry over 23 days, through the stages of lytic infection and recovery (Fig. 1b). Uninfected *E. huxleyi* cultures reached cell density of ∼5 × 10^6^ cells mL^-1^ within 5 days and remained steady until day 23. In contrast, the virus-infected cultures experienced a profound decline during days 2-7 post-infection, whereby only < 0.5% of the culture survived (Fig. 1b). The fraction of dying cells was measured by Sytox Green staining and reached 80% by day 4 (Fig. S1). Cultures remained at a minimal cell density of ∼10^3^ cells mL^−1^ for four days. From day 11, cell abundance increased, signifying the recovery phase of the cultures. Recovery dynamics varied between the biological replicates (n=3), but eventually, all recovered. Replicate A steadily grew and reached the starting point concentration on day 21, but replicates B and C fully recovered only on day 37 (Fig. S2). A divergence between the replicates during the recovery phase is expected, given that we are comparing processes that occur after almost complete lysis of the culture. Deviation between the replicates in the number of surviving cells, which are the foundation of the new resistant population, can lead to less synchronization in the recovery dynamics. After the last sampling point (day 23), the three recovered cultures were diluted into fresh media and kept under regular growth conditions.

The onset of virus release (extracellular virions) was already apparent at 8h post-infection, and virus abundances increased over 100-fold within the first two days (Fig. 1b). This preceded the significant rise in cell death, suggesting viruses were released by budding (Vincent *et al*., 2021). A constant high viral particle load, over 10^8^ viral particles mL^-1^, was measured through the recovery of host cultures, suggesting the cells developed resistance to EhVM1 and transitioned to a co-existence state (Joffe *et al*., 2024). To validate the acquisition of resistance to viral infection in the recovered host cultures, we challenged them with a fresh EhVM1. Cell abundance of the challenged recovered cultures was similar to uninfected controls (Fig. 1c and S3), meaning the recovered cultures gained resistance to EhVM1.

### The transcriptomic response of *E. huxleyi* to viral infection

To understand the phenotypic switch from sensitivity to resistance to viral infection, we conducted a detailed dual host-virus transcriptomic analysis during the 23-day time series. Following the same cultures of one evolving strain from sensitive to resistant, we aimed to identify the genes contributing to the resistance phenotype. We collected RNA samples from 9 time points (0.5h; 1-4, 7, 15, 19, 23 days post-infection) throughout the time series, with an additional sampling point on day 50, after the recovered cultures grew in fresh media (Fig. 1b).

In total, 48 samples underwent dual host-virus 3’ RNA sequencing (RNA-seq) using the MARS-seq protocol (Keren-Shaul *et al*., 2019), and reads were mapped to a pan-transcriptome reference containing both *E. huxleyi* and EhV (Feldmesser *et al*., 2021; Fromm *et al*., 2022). The dual RNA-seq captures both host and virus gene expression and enables linking the host transcriptional state with the progression of infection. To profile the transcriptional signature of infected and resistant cells, we focused on the samples from the infection and recovery phases. We included time points from 0.5h post-infection (T0), and days 1, 2, 3, 19, 23, and 50 post-infection (T1, T2, T3, T19, T23, T50). Respectively, the relevant time points from uninfected samples were included (T0 to T3). Only genes with more than 10 read counts in at least three samples were included in the transcriptomic analysis, yielding 19,816 expressed host genes and 270 virus genes (Supporting Data S1). The number of expressed *E. huxleyi* and EhV genes aligns with findings from previous transcriptomic studies (Rosenwasser *et al*., 2014; Ku *et al*., 2020; Feldmesser *et al*., 2021; Bousquet *et al*., 2025).

Principal component analysis (PCA) of host gene expression revealed distinct transcriptional profiles of recovered and infected populations. PC1, which accounted for 46.7% of the variance (Fig. 2a), separated the recovery phase from the uninfected and infected phases. The recovery phase samples (T19, T23, T50) are distributed along the PC1 axis but do not form time-specific clusters, similar to the observed asynchrony in recovery dynamics (Fig. 1b). PC2 accounted for 22.3% of the variance and captured the temporal progression in the infection phase, based only on the host response to infection. Uninfected samples clustered closely, displaying relatively low variation throughout the sampling period. Together, these results highlight the transcriptional plasticity during infection and recovery phases.

**Figure 2.**
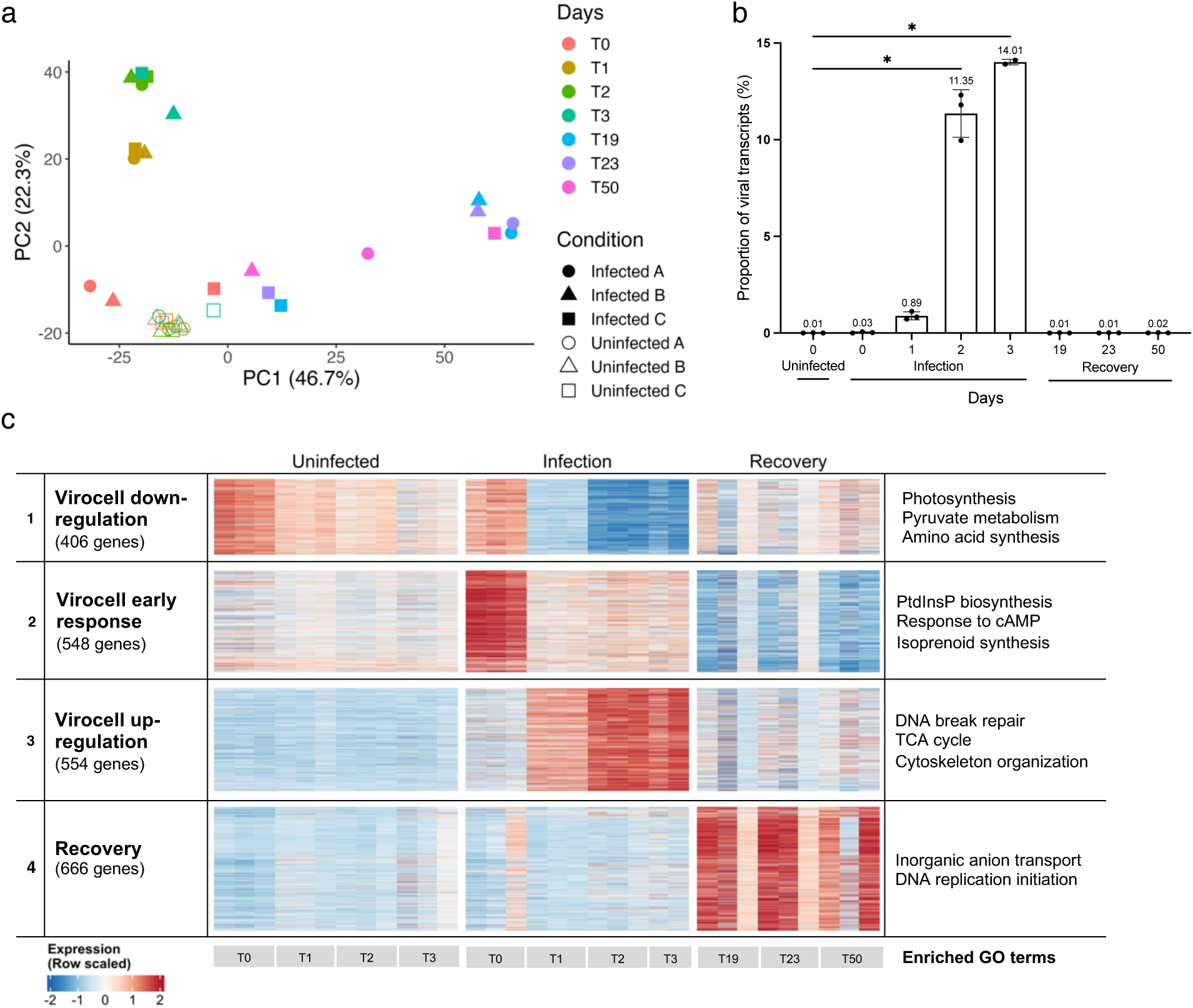
Distinct transcriptional patterns of *E. huxleyi* during infection and recovery. (a) PCA based on host gene expression values (normalized counts after variance stabilizing transformation, VST) separated the population into the different states of infection and recovery. Colors represent time and shape the condition and replicate. (b) The proportion of viral RNA transcripts out of the total transcripts increased with the progression of viral infection. Asterisks represent a statistically significant increase (one-way repeated measures ANOVA and Dunnett’s post-hoc multiple comparison tests, p < 0.05). (c) Expression patterns of the 2,174 significantly differentially expressed host genes during EhV infection and recovery, clustered with K-means (k=4). The heatmap shows expression patterns (z-scores VST values) of each gene (row) along the different replicates of each timepoint (column). To the left are cluster names and the total number of genes in each cluster. To the right are significantly enriched topGO terms (Fisher test, biological process ontology, p-values ≤ 0.01). Abbreviations: PtdInsP - phosphatidylinositol phosphate.

The proportion of viral RNA transcripts out of the total mapped transcripts increased from ∼0.03% at T0 to 14% at T3 (Fig. 2b). Previous studies on other EhV strains detected a near-complete takeover of the host’s transcriptional machinery and mRNA pool, reaching 80% of the reads within 24 hours (Rosenwasser *et al*., 2014; Ku *et al*., 2020). The lack of viral takeover on the global transcriptome here offers an opportunity to study the previously obscured transcriptional response of *E. huxleyi* throughout infection.

Gene expression analysis along the time series revealed 2,174 DE host genes in at least one comparison (absolute log2(Fold Change) ≥ 1, FDR p-value < 0.05; Supporting Data S2). We conducted the DE analysis on *E. huxleyi* genes only since introducing another organism (virus) into a finite pool of reads can generate false identification of DE genes, as RNA-seq is a compositional data representing relative abundances of genes.

Unsupervised k-means clustering (k=4) highlighted temporal expression patterns with significant enrichment in specific GO terms (Fig. 2c) and KEGG pathways (Supporting Data S3). The host response to infection is represented by clusters 1-3 and includes an early response detected after 0.5h (T0) and an extensive reprogramming of the virocell transcriptome at T1-T3. The virocell early response cluster (cluster 2; 548 genes) captured an immediate and transient gene up-regulation, marked by genes involved in cellular signaling pathways and the synthesis of signaling molecules (Fig. S4). Among them are phosphatidylinositol (PtdIns) kinases, ras family GTPases, lipoxygenases producing oxylipins, and isoprenoid synthesis genes. PI3K, a PtdIns kinase, is known to be activated as a stress-induced response in higher eukaryotes (Munnik *et al*., 1998; Ramanan *et al*., 2018), and oxylipins are released in diatoms in response to viral infection (Edwards *et al*., 2024), grazing stress, or disrupted membranes. These oxylipins act as a chemical signal to induce programmed cell death in neighboring diatom cells (Vardi *et al*., 2006). Additionally, genes encoding proteins with domains homologous to Toll/interleukin-1 receptor/resistance (TIR) domains, a signaling component of the eukaryotic immunity system, were detected.

Virocell clusters 1 and 3 reflect a global remodeling of core metabolic pathways to support viral propagation, in corroboration with previous findings (Rosenwasser *et al*., 2014) (Table S1). The virocell down-regulated cluster (cluster 1; 406 genes) was enriched with pathways associated with core cellular processes such as photosynthesis, amino acid metabolism, pyruvate metabolism, and terpenoid synthesis via the 2-*C*-methyl-d-erythritol-4-phosphate (MEP) pathway. The virocell up-regulated cluster (cluster 3; 554 genes) involved pathways of DNA break repair and DNA replication, TCA cycle enzymes, and cytoskeleton organization.

The recovery cluster (cluster 4; 666 genes) grouped genes that were upregulated during the recovery phase. It highlighted a unique transcriptional program of the cells that acquired resistance, distinct from the gene expression patterns of uninfected or infected cells. This cluster was enriched with inorganic anion transmembrane transport pathways and DNA replication initiation (Table 1).

**Table 1.**
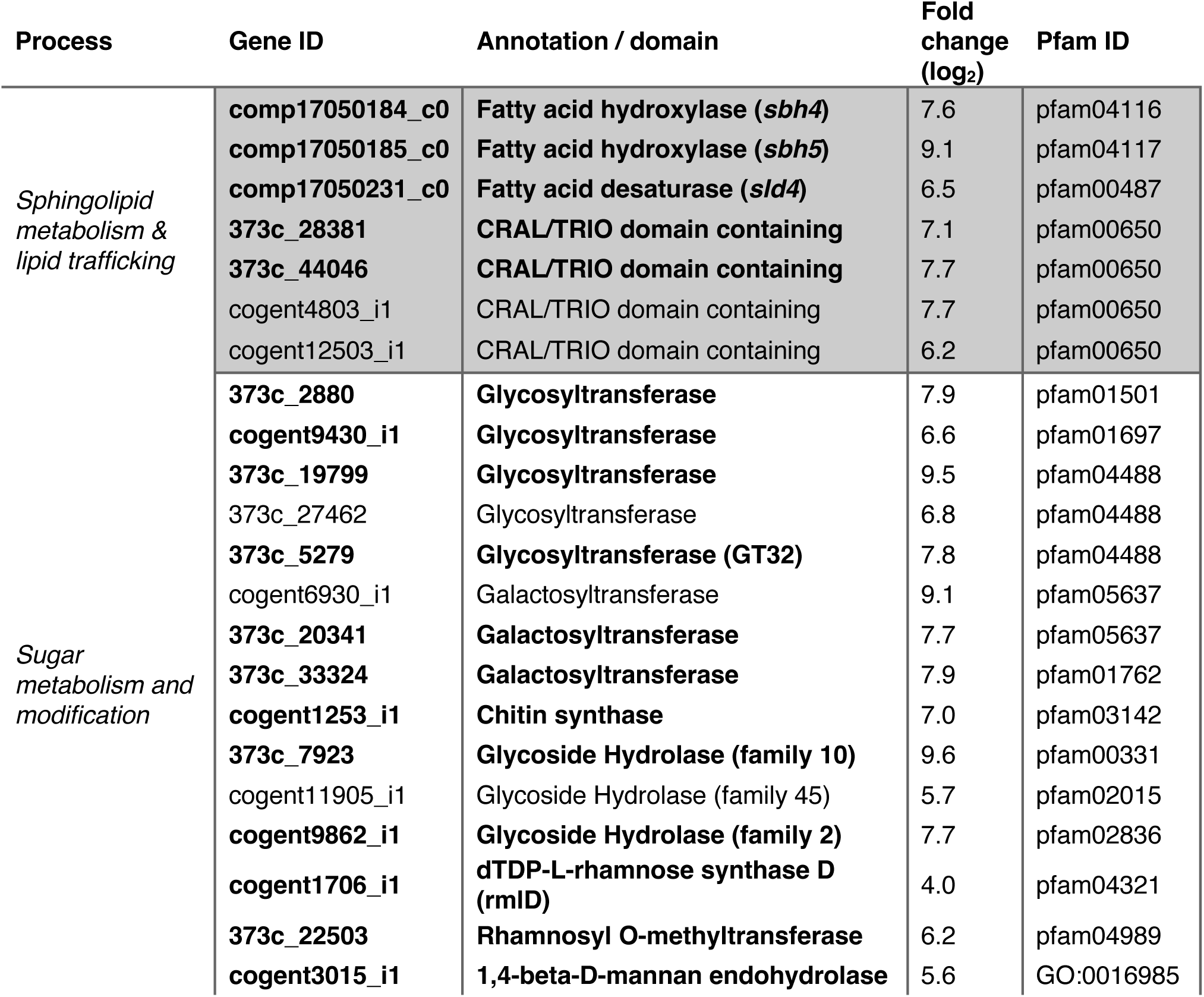
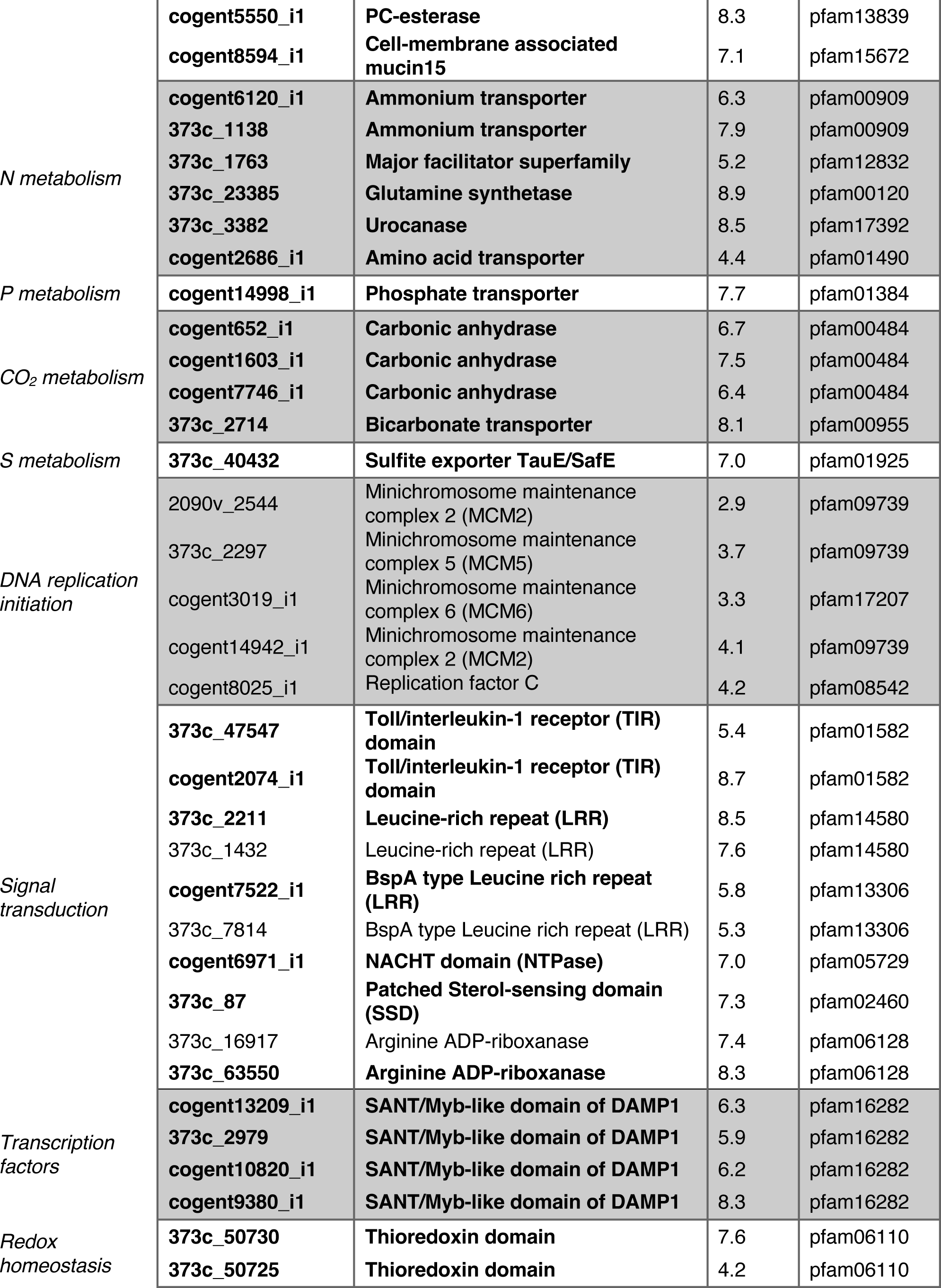
Overview of up-regulated functions in resistant *E. huxleyi* populations. Some functions are represented by multiple genes (Supporting Data S4). GO terms were added when Pfam was missing. Fold change values represent differential expression between T19-infected samples and T0-uninfected samples (log2). Lines in bold represent genes that are upregulated in all resistant strains based on the comparative transcriptomics analysis.

### The unique transcriptional state of recovered cells

To validate the transcriptional profile of recovered cells that gained resistance, we repeated the experiment and infected a fresh culture of *E. huxleyi* RCC6946 with EhVM1. The resistant population that recovered from this infection was kept under regular growth conditions for several weeks. Then, the cells were sampled for RNA-seq and compared to the sensitive ancestor strain at a single time point during the exponential growth phase. The analysis resulted in 787 DE genes (absolute log2(FC) ≥ 1, FDR p-value < 0.05). Approximately half of the detected DE genes (391/787) were also DE in the time series transcriptome and were mainly from the recovery cluster (351 genes). We focused on the genes shared between the recovery phase of the two independent experiments, as they have a stronger association with the resistant phenotypic state.

Among the shared genes, we detected the three lipid modification enzymes predicted to generate specific lipid markers of resistant cells (resGSLs) (Schleyer *et al*., 2023) (Table 1). These include sphingolipid desaturase 4 (*sld*4), sphingoid base hydroxylases 4 and 5 (*sbh*4-5), and several CRAL/TRIO domain-containing genes with putative roles in lipid intracellular trafficking (Panagabko *et al*., 2003). Sphingolipids (SLs) belong to a lipid class with important signaling and membrane structure roles in all eukaryotes and some bacterial taxa (Aguilera-Romero *et al*., 2014). Altering the sphingolipid composition may provide resistance by activating cellular signaling pathways or by directly preventing viral adsorption and entry to the cells through membranal lipid raft remodeling, the possible mediators of viral entry and exit (Mackinder *et al*., 2009; Rose *et al*., 2014).

Altering glycan structures on the host cell surface can also influence host-virus interactions. Membrane-bound glycans (glycolipids and glycoproteins) are present on both the host cell surface and viral particles. Viruses utilize these glycans for attachment and entry into the cell. In the recovered cells, sugar modification enzymes that can remodel membrane-bound glycans were found to be unregulated. These include glycosyltransferases (GT), sugar methyltransferases, and glycoside hydrolases (GH) that can modify cell surface polysaccharides (Table 1). Specifically, xylanase and cellulase enzymes from GH families specializing in cell wall polysaccharide degradation were upregulated (GH5, GH10, determined by the CAZy Database (Drula *et al*., 2022)). Also, some of the GT and GH families were predicted to localize extracellularly, according to Deeploc (Ødum *et al*., 2024), potentially modifying cell surface glycans. Similarly, in the green alga *Ostreococcus tauri*, resistant cell lines that recovered from infection by a giant virus (OtV) overexpress numerous glycosyltransferases (Yau *et al*., 2016). Additionally, giant viruses (NCLDV) are known to encode enzymes for polysaccharide manipulation and synthesis of sugar-nucleotides and glycans (Speciale *et al*., 2022). In particular, chloroviruses can remodel the cell walls of their host during infection. They encode GTs that synthesize the glycans attached to viral major capsid glycoproteins (DeAngelis *et al*., 1997; Van Etten *et al*., 2017). EhV itself encodes for multiple GTs and GHs, suggesting their significance in the viral life cycle. Thus, upregulating enzymes to manipulate cell surface glycans could potentially contribute to host resistance.

In the resistant cells, we also detected higher transcription of genes involved in nutrient uptake and metabolism, mainly nitrogen (N), phosphorus (P), CO2, and sulfate (S). Among these are ammonium transporters, glutamine synthetase, urocanase, phosphate transporters, sulfite exporters, carbonic anhydrase, and bicarbonate transporters. Upregulating transcription of nutrient uptake genes usually reflects nutrient-limited cells (Rokitta *et al*., 2014; Alexander *et al*., 2020). Under nutrient limitation, cells induce high-affinity uptake systems and allocate nutrients from internal storage to handle the scarcity, which might result in a transient resistant state. For example, under P stress, *E. huxleyi* cells exhibit membrane remodeling by substituting membrane phospholipids with non-phosphorus lipids (Van Mooy *et al*., 2009; Shemi *et al*., 2016). EhV infection under P stress resulted in lower viral production, without apparent bloom demise (Bratbak *et al*., 1993). Additionally, we detected the expression of SANT/myb transcription factors (TFs). Myb TFs are abundant in plants and microalgae (Thiriet-Rupert *et al*., 2016) and are involved in regulating nutrient stress responses to P, N, and CO2 in several microalgae lineages (Yoshioka *et al*., 2004; Kumar Sharma *et al*., 2020; Wang *et al*., 2022). In *E. huxleyi*, specific myb TFs are expressed in resistant haploid cells (Von Dassow *et al*., 2009) and during the recovery of resistant cells (Frada *et al*., 2017). Thus, the SANT/myb TFs detected here might be involved in regulating the expression of nutrient-stress genes in *E. huxleyi* resistant cells.

Interestingly, among the upregulated genes in the recovered-resistant population were genes potentially involved in pathogen-recognition signaling pathways. The proteins encoded by these genes contain motifs or domains that are conserved in eukaryotes and activate an innate immune response upon pathogen recognition. The immune response typically includes activation of transcription factors that initiate the transcription of immunity genes. Here, we observe that the recovered cells highly transcribe genes encoding Toll/interleukin-1 receptor/resistance (TIR) domains, leucine-rich repeat (LRR) motifs, and NACHT domains (NTPase), which have functional homologs involved in the immune responses of eukaryotic cells. A potential outcome of this signal transduction is an unfavorable environment for the viral replication cycle. Taken together, recovery from viral infection involves several key processes, including signaling pathways associated with innate immune responses, remodeling of lipids and membrane glycans, and nutrient uptake and metabolism. It remains to be determined if the pathways discussed above serve as the molecular mechanisms that provide resistance or if they indicate a unique metabolic state of recovered cells. For instance, the expression of nutrient-stress genes, even in the presence of adequate growth conditions, can arise from an induced response that actively prevents the virus from utilizing them. Alternatively, it could be a secondary effect caused by changes in the cell membrane that block viral entry but also impact nutrient uptake. Indeed, resistant cells tend to carry a fitness cost, which is reflected by a slower growth rate or a reduced carrying capacity (Joffe *et al*., 2024). Due to the lack of genetic tools in *E. huxleyi*, these questions remain open for future research aiming to genetically manipulate these gene candidates and examine changes in susceptibility to viral infection.

### Resistant *E. huxleyi* strains share a conserved transcriptional state

Next, we asked whether the transcriptional state of recovered populations is similar to that of other resistant *E. huxleyi* strains. We performed a meta-analysis comparing the gene expression profiles of multiple strains that exhibit resistance to different EhVs. These strains were either isolated as resistant against all tested EhVs or recovered from EhV infections. With this approach, we aimed to identify a core set of genes associated with resistance to viral infection. This could provide insights into the molecular mechanism underlying resistance and serve as potential biomarkers for the specific identification of resistant *E. huxleyi* cells.

We compared the DE genes identified in this study with expression patterns from available published transcriptomes of resistant and sensitive *E. huxleyi* strains (see Materials and Methods). Among the 2,174 DE genes in the time course dataset, we identified those significantly DE (upregulated or downregulated) also in the other resistant strains. These shared DE genes were then grouped based on the cluster affiliation from Fig. 2c (Supporting Data S5).

Our analysis revealed that most genes in the recovery cluster were also upregulated in other resistant strains (Fig. 3a, height of the bars, cluster 4). In contrast, the virocell clusters (clusters 1-3), representing a response to infection, had a lower degree of similarity with the transcriptome of resistant strains. The upregulated genes in resistant strains also showed higher fold change (FC) values, with expression levels ranging from 2^5^ to 2^15^ times higher than those in sensitive strains. The extreme FC values suggest these genes are not expressed by the sensitive strains (Fig. 3b). These findings strengthen the functional importance of genes from the recovery cluster in the phenotypic state of resistance.

**Figure 3.**
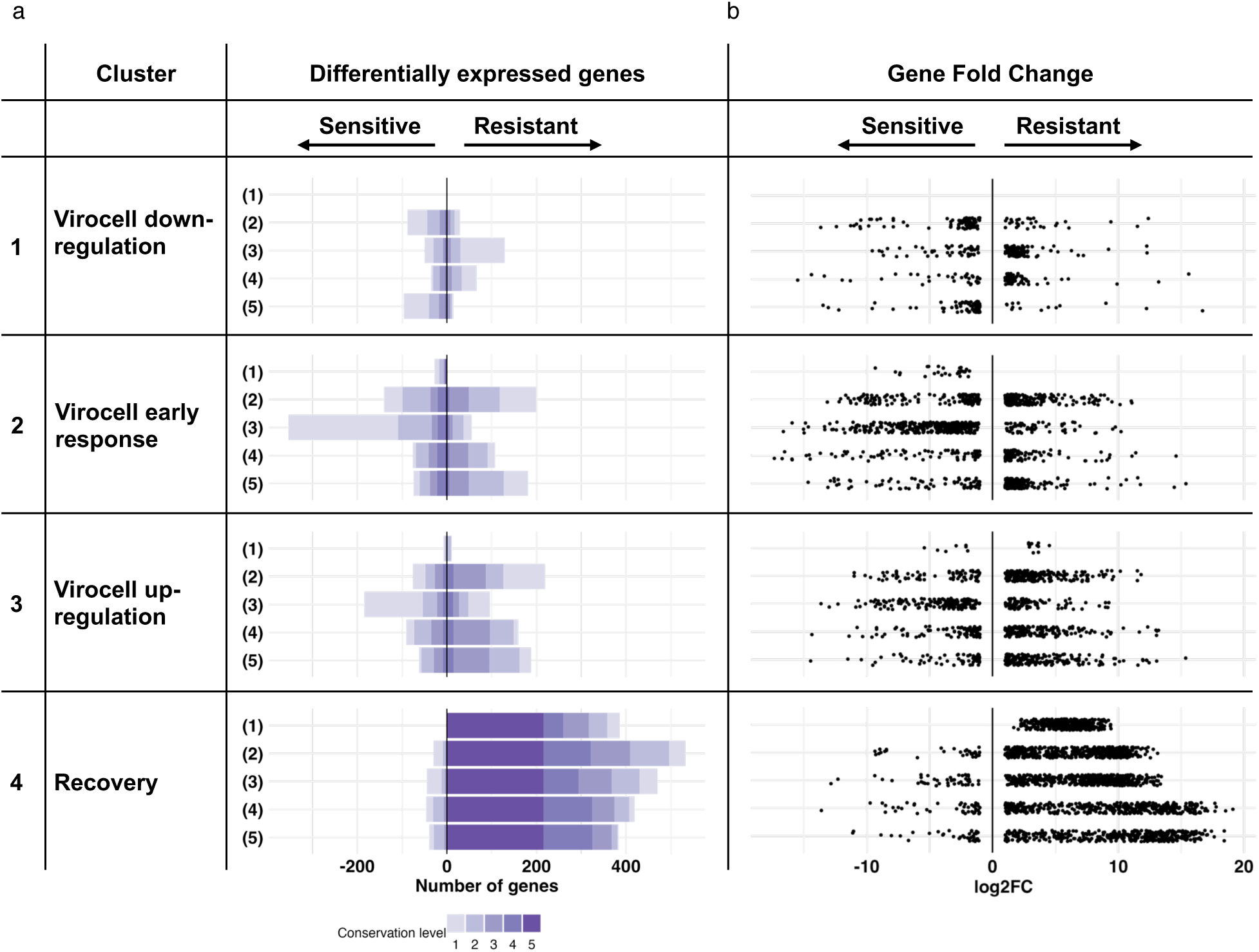
Expression patterns in transcriptomes of resistant *E. huxleyi* strains. (a) Differentially expressed genes in each resistant vs. sensitive strain transcriptome comparison, organized by the cluster affiliation of Fig. 2c. The x-axis counts significantly differentially expressed genes in each transcriptome (|log2(FC)| > 1, FDR p-value < 0.05). Arrows indicate the direction of differential expression: a positive count for genes upregulated in the resistant strain, and a negative count for genes downregulated in the resistant strain (i.e., upregulated in the sensitive strain). The conservation level (color gradient) reflects the number of transcriptomes in which a gene was upregulated or downregulated. (b) Distribution of fold change (FC) values (log2) of the differentially expressed genes from (a). Transcriptome numbering refers to the resistant strains as follows: (1) *E. huxleyi* RCC6946 recovered population from EhVM1, results from this paper; (2) *E. huxleyi* CCMP373 from Feldmesser *et al*., 2021; (3) *E. huxleyi* CCMP379 from Kendrick *et al*., 2014; (4) *E. huxleyi* LC5-11A (1N-flagellated) and (5) LC5-12A (2N-flagellated) from Bousquet *et al*. 2025.

Next, we assessed if the DE genes from each transcriptome formed a mutually shared group across the transcriptomes. We assigned each gene a score ranging from 1 to 5, reflecting the number of resistant strains in which it was upregulated, serving as a conservation level across the resistant phenotype. Similarly, downregulated genes received a score from -1 to -5 based on the number of transcriptomes (Fig. 3a, color gradient). We identified a core group of 206 genes (“resistance genes”) within the recovery cluster that were upregulated in all the tested resistant strains (conservation score=5). This conserved group of genes represents a potential molecular signature of *E. huxleyi* resistance.

The core resistance group covers most of the genes discussed above, mainly the immune signaling pathways, glycan and lipid remodeling and metabolism, and nutrient uptake (Table 1, bolded genes). Additionally, using AlphaFold 3 (Abramson *et al*., 2024) we generated structural predictions for a better functional annotation of genes with no sequence-based homologs. This revealed new functional groups, highlighted by genes involved in RNA/DNA processing and calcium (Ca^2+^) binding and transport (Supporting Data S6). Among them are genes with zinc-finger domains, EF calcium-binding motifs, and bestrophins, which function as anion channels in humans. Interestingly, the only process not conserved with other resistant strains is the DNA replication initiation (Table 1), suggesting it is a unique transcriptional signature of recovery in the *E. huxleyi* RCC6946 strain.

We also assessed whether the expression of the resistance genes is conserved with other eukaryotic microalgae, but the major challenge is the lack of available transcriptomes for resistant isolates. The one available transcriptome of resistant *O. tauri* cell lines (Yau *et al*., 2016) revealed homologs to two resistance genes (ostta19g00630, ostta19g00120) that are highly expressed in a resistant *O. tauri* isolate. The two genes are predicted to be sugar modification enzymes – galactosyltransferase and glycosyltransferase, respectively. This further supports that glycan modifications are a common metabolic strategy to alter susceptibility to viral infection.

Taken together, cells that develop resistance to viral infection exhibited expression patterns that were conserved across multiple resistant strains of *E. huxleyi*. The resistance genes highlight a specific profile of resistant cells. This profile includes genes that influence the carbohydrate and lipid composition of the cells, as well as the metabolism of macronutrients. Significantly, TIR-containing genes—known immune response components that are evolutionarily conserved across plants and animals—are reported here for the first time as being involved in the immune response of algal cells.

### Exploring the functional role of TIR domains in algal resistance

Due to the principal role of TIR domain-containing proteins in the immunity of animals, plants, bacteria, and archaea (Essuman *et al*., 2017; Fitzgerald & Kagan, 2020), we focused on elucidating their transcriptional patterns during alga-virus dynamics. TIR domain-containing proteins have diverse functions, acting as immune signaling mediators or degradation enzymes. Recently, studies have shown that TIR domains are involved in bacterial immune response by recognizing phage infections (Ofir *et al*., 2021). Upon infection, a TIR-based signaling pathway activates an effector protein that depletes cellular nicotinamide(Tang *et al*., 2019) adenine dinucleotide (NAD^+^), leading to cell death that aborts phage replication (Ka *et al*., 2020; Ofir *et al*., 2021; Tamulaitiene *et al*., 2024). In the context of the ongoing evolutionary host-virus arms race, other studies suggest that phages can overcome bacterial defense by sequestering the TIR-derived signaling molecule (Leavitt *et al*., 2022) or by restoring NAD^+^ levels in infected cells (Osterman *et al*., 2024).

Microalgae and macroalgae (multicellular algae) share immune motifs that are common in both plants and animals, such as TIR, nucleotide-binding site (NBS), and leucine-rich repeat (LRR) domains (Richter & Levin, 2019; Kloareg *et al*., 2021). In the red marcoalga *Pyropia yezoensis*, the transcriptional response to infection by the pathogen *Pythium porphyrae* involves upregulation of genes with LRR, NBS, and TIR domains (Tang *et al*., 2019). However, to date, TIR-based anti-viral systems have not been described in the evolutionary branch of microalgae. Therefore, we searched for all TIR-containing genes encoded by *E. huxleyi*. Based on conserved domains search, we found 15 genes predicted to encode a TIR domain which were DE during the time series transcriptome (Fig. 4a). The TIR domain-containing genes exhibit temporal expression patterns, some upregulated exclusively in the recovered phase (T19-T50), some immediately after viral inoculation, as part of the virocell early response phase (T0) and some transcribed mainly during the infection (T1-T3). The temporal expression in various phases of infection and recovery may suggest different functions, e.g., signal transduction or NAD^+^ hydrolysis.

**Figure 4.**
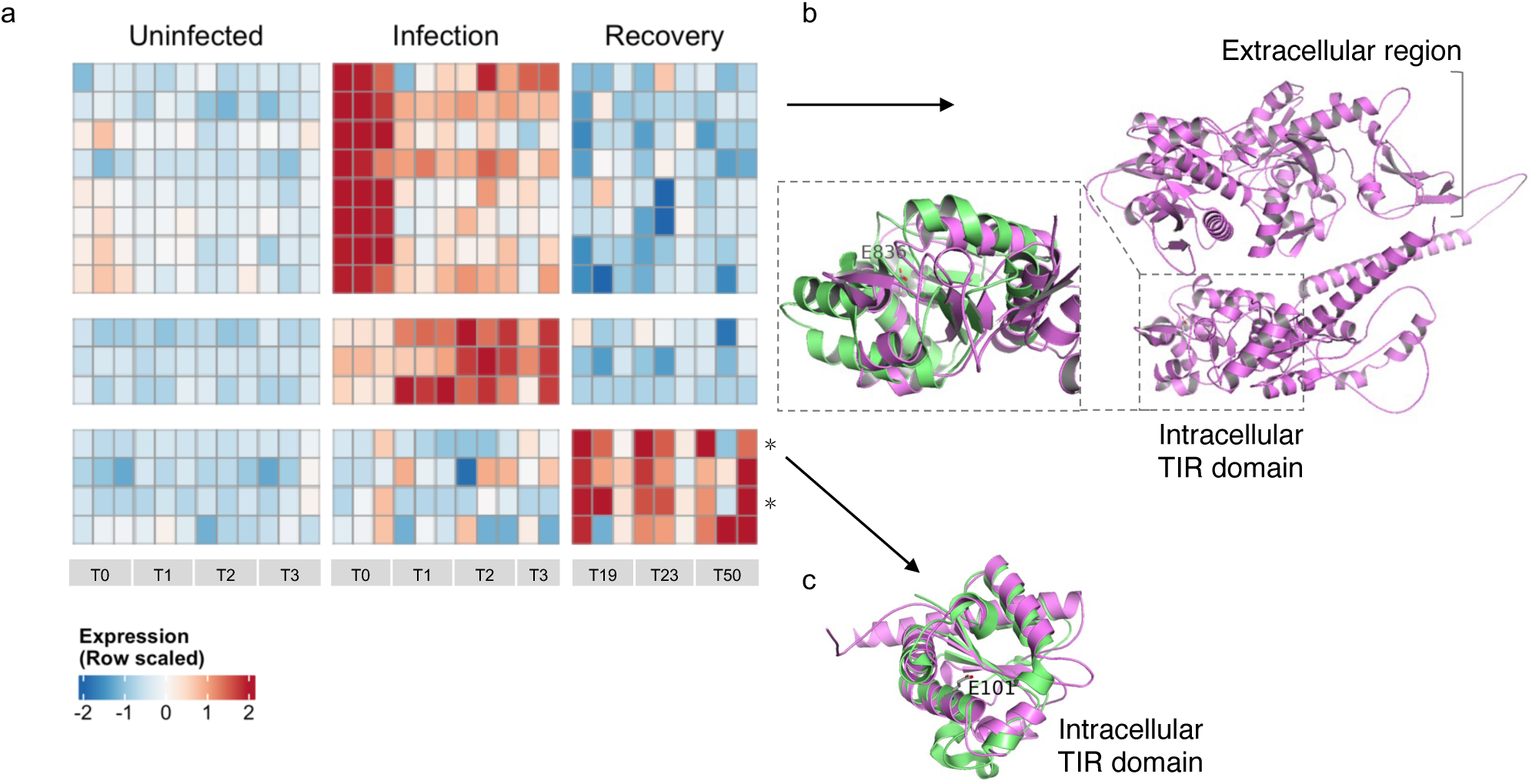
Expression patterns of TIR domain-containing genes. (a) The heatmap shows expression patterns (z-scores VST values) of each gene (rows) along the different replicates of each timepoint (columns). Cluster numbers are to the left. Asterisks mark the two genes expressed by all resistant strains. (b) *E. huxleyi* cogent1101_i1 structural prediction (violet) reveals an extracellular region followed by an intracellular predicted TIR domain with the conserved glutamic acid residue (E836). In the highlighted box is the predicted TIR domain, aligned with the crystal structure of the *Arabidopsis thaliana* TIR domain of disease resistance protein RPS4 (4C6R; green). (c) *E. huxleyi* 373c_47547 structural prediction (violet) shows a predicted intracellular TIR domain with the conserved glutamic acid residue (E101). The domain is aligned with the crystal structure of the *Homo Sapiens* TIR domain in SARM1 bound to an NB-7-ADPR adduct (8D0D; green). Only the domain is shown since AlphaFold could not accurately predict a structure for the rest of the protein.

To characterize the expressed TIR domain-containing genes, we generated structural predictions using AlphaFold 3 (Abramson *et al*., 2024). Typically, TIR domain-containing genes are associated with other domains, such as NBS and LRR. However, the TIR domain-containing genes in *E. huxleyi* did not encode such additional domains. It’s worth noting that there are known TIR domain-containing genes that possess only the TIR domain (“truncated” TIR) and still function as immune receptors (Nishimura *et al*., 2017; Santamaría *et al*., 2019; Tian *et al*., 2021; Ogden *et al*., 2023). Importantly, while the length of the detected TIR domains varied (ranging from 76 to 128 amino acids) and their sequences differed, the putative catalytic glutamic acid residue in the active site (Essuman *et al*., 2017; Wan *et al*., 2019) was conserved (Fig. S5, Supporting Data S7).

Additionally, *E. huxleyi* TIR domains show structural similarity to known, functionally active TIR domains found in bacteria, plants, and humans. To assess whether the temporal expression patterns (Fig. 4a) imply different functions, we examined shared features among the cluster members, such as additional encoded domains and predicted transmembrane regions. We found that TIR domain-containing genes in the virocell clusters (clusters 2 and 3) were predicted to have an extracellular region or multiple transmembrane helices at the N-terminus. Following these regions was a TIR domain, with structural homology to signal-producing TIRs. For example, the cogent1101_i1 gene from cluster 2 consists of three main components: an extracellular region, a transmembrane helix, and an intracellular TIR domain (Fig. 4b). The extracellular region showed structural similarities to ligand-binding domains, such as those found in leucine-binding and epidermal growth factor (EGF)-like domains. The TIR domain shared structural homology to signal-producing TIRs, like the TIR domain from *Arabidopsis thaliana* disease resistance protein, RPS4. Furthermore, the early expression (0.5h post-infection) of TIR domain-containing genes from cluster 2 resembles the early activation of TIR signaling in plants in response to pathogen-associated molecular patterns (PAMP) (Bjornson *et al*., 2021; Tian *et al*., 2021). This provides initial evidence that TIR domain-containing genes from the virocell clusters may have a receptor-like function to generate signaling molecules in response to viral infection.

The TIR domain-containing genes from the recovery cluster — particularly the two expressed by all resistant strains (Fig. 4a, marked by asterisks) — were predicted to localize intracellularly in the cytoplasm (Supporting Data S7). These two genes (373c_47547, cogent2074_i1) show high structural homology to signal-producing TIRs and the Human SARM1 TIR domain (Fig. 4b). SARM1 functions as an NAD^+^ hydrolyzing enzyme that triggers cell death and neuronal degradation, effectively halting the progression of viral infections (Sundaramoorthy *et al*., 2020). When cells activate SARM1, they can restrict the spread of viruses and protect neighboring cells from infection; however, this protection comes at the cost of their own death. In other words, the expression of TIRs prevents the infection but does not save the cell itself.

However, resistant *E. huxleyi* cells express TIR domain-containing genes without experiencing cell death, even when challenged with EhV. One possible explanation for this phenomenon is that the TIR domain-containing genes act only as a secondary line of defense. According to this hypothesis, resistant cells first try to prevent infection through morphological changes, such as glycan and lipid modifications. But, should this strategy fail, the cells also pre-activate transcription of TIRs in a self-inhibition state to prevent unnecessary cell death. If a virus successfully invades the cell, those TIRs will be activated, leading to cell death and inhibition of viral spread within the population. Another possible explanation is that these TIRs do not cause cell death; they may function as transcriptional regulators of stress responses. Research in plants has shown that adding 2’,3’-cyclic adenosine monophosphate (2’,3’-cAMP), which is synthesized by TIR proteins (Yu *et al*., 2022), can activate transcriptional reprogramming related to stress responses (Chodasiewicz *et al*., 2022).

Interestingly, TIR domains are relatively rare in viral genomes (Spear *et al*., 2009; Iyer *et al*., 2022). However, several NCLDVs that infect eukaryotic microalgae from the haptophyta phylum do encode TIR domain-containing genes (Gallot-Lavallée *et al*., 2015; Bhattacharjee *et al*., 2023; Wang *et al*., 2025). Among these are *Haptolina ericina* virus (GenBank: XPO57143.1), *Chrysochromulina ericina* virus (ALH23162.1), *Prymnesium kappa* virus (XPO57813.1), and an unidentified species within the Nucleocytoviricota phylum (UZT28792.1). It is known that bacterial pathogens of plants produce bacterial TIR-like virulence effectors that resemble plant-derived TIR products (Eastman *et al*., 2022; Hulin *et al*., 2023). These effectors could negatively regulate host immunity or manipulate NAD^+^ metabolism. Therefore, the function of viral-encoded TIRs in marine NCLDVs is still to be explored, but it has the potential to influence host immune pathways.

## Conclusions

Our study provides a detailed mapping of the transcriptional landscape during the transition from algal sensitivity to resistance in response to viral infection. At the cellular level, viral infection triggers a transcriptome remodeling of key pathways, likely supporting viral replication. At the population level, this infection imposes selective pressure, revealing the transcriptional profile associated with the resistant phenotype. This transcriptional profile was consistently observed across various resistant strains of *E. huxleyi*, indicating a core resistance expression program. Some of these immunity elements, such as the TIR domains, were evolutionarily conserved through branches in the tree of life. Nonetheless, functional assays are needed to elucidate the mechanisms underlying the resistance phenotype.

The transcriptional landscape of resistance in *E. huxleyi* against a giant virus infection could provide valuable insights into how microalgae defend themselves against giant viruses. Various haptophytes and other marine microalgae are frequently infected by giant viruses (NCLDV). But, currently, the molecular basis for resistance mechanisms against giant viruses remains largely unknown. The group of resistance-related genes revealed here is composed of those involved in signaling pathways associated with innate immune responses (such as TIR domains), remodeling of lipids and membrane glycans, as well as nutrient uptake and metabolism. Therefore, sequencing the transcriptomes of additional resistant strains may help determine whether these transcriptional changes are common elements of resistance against giant virus infection among different microalgae species.

The discovery of core resistance-related genes can also be used to study how resistance evolves within a population. Do resistant cells exist before viral exposure, persisting as a small subpopulation due to the associated costs of resistance? Alternatively, is resistance induced in infected cells that abort the infection cycle or in uninfected bystander cells as a response to chemical cues released by infected cells? Applying techniques such as dual single-cell RNA-sequencing (scRNA-seq) will enable sensitive detection of resistant cells using the core resistance genes. Then, the simultaneous view of the viral transcriptome at the individual cell level can provide valuable insights into the origins and development of resistant cells in the population.

Additionally, the group of core resistance-related genes represents potential gene biomarkers for detecting resistant *E. huxleyi* cells in natural oceanic blooms. This enables the quantification of resistant cells in host-virus dynamics during algal blooms. Resistant cells may have a significant ecological role in bloom succession as the survivors of viral infections and the seed population of the next year bloom. Resistant cells may also impact carbon export to the deep sea through cell aggregation. The resistance genes include several genes involved in glycan modifications and the degradation of polysaccharides. Production of extracellular acidic polysaccharides, called Transparent Exopolymer Particles (TEP), is associated with particle aggregation (marine snow). During viral infections, the production of TEP increases, which enhances particle aggregation (Vardi *et al*., 2012; Laber *et al*., 2018; Vincent *et al*., 2021) and carbon export to the deep sea through a process known as the “virus shuttle” (Sullivan *et al*., 2017). Thus, aggregation of resistant cells can contribute to carbon export in a parallel mechanism. Furthermore, environmental factors, such as increased temperatures, have been shown to induce a resistant state in *E. huxleyi* cells (Kendrick *et al*., 2014). Consequently, detecting resistant cells using the newly identified gene biomarkers, can be of great value in assessing the impact of elevated temperature on host-virus dynamics in light of climate change.

## Acknowledgments

This study was supported by the Simons Foundation grant (no. 150809), “Elucidating the unknown mechanisms of viral resistance in algal blooms in the ocean” awarded to AV.

## Author contributions

TS and AV planned and designed the project. TS and DS performed the experiments and prepared transcriptomics libraries. SBD and EF constructed the reference transcriptome. TS and AF performed gene annotation and meta-analysis. TS conducted all other data analyses. TS and AV wrote the manuscript.

## Data availability

Data are available in the main text or supporting information. The raw sequence reads for the two transcriptomes analyzed in this paper were deposited on NCBI and are pending accession. The *E. huxleyi* reference transcriptome (pan-transcriptome) files, including transcripts and protein fasta files, as well as GO and KEGG functional annotation files, are available via the Figshare repository: doi.org/10.6084/m9.figshare.28891220.

## Materials and Methods

### *E. huxleyi* and EhV strains

The *E. huxleyi* strain RCC6946 and the EhV strain EhVM1 were isolated from an *E. huxleyi* bloom during a mesocosm experiment in Raunefjorden, Bergen, Norway (60.27°N; 5.22°E) in May 2018.

### Culture growth conditions and experimental setup

*E. huxleyi* RCC6946 cultures were grown in filtered autoclaved seawater (FSW) supplemented with a modified K/2 medium (Keller *et al*., 1987) (18-μM KH2PO4 instead of organic phosphate), and the antibiotics Ampicillin (100 μg mL^-1^) and Kanamycin (50 μg mL^-1^). Cultures were incubated at 18°C with a 16:8 h light-dark cycle and 100 μmol photons m^−2^ s^−1^ light intensity provided by white light-emitting diodes. EhVM1 was propagated on *E. huxleyi* RCC6946 until the algal culture lysed to clearance. The viral lysate was filtered through a 0.45μm pore size Nalgene Rapid-Flow filter unit (PES, Thermo Fisher Scientific) and stored in the dark at 4°C until the infection assay. In total, we performed two infection assays. In the time series infection assay, exponential phase RCC6946 cultures (∼10^6^ cells mL^-1^, volume of 1.4L) were infected with EhVM1 in a 0.075:1 virus-to-cell ratio (Multiplicity of Infection; MOI). Since a high MOI (5:1) triggers a rapid shutdown of most host genes (Rosenwasser *et al*., 2014; Ku *et al*., 2020), using a low MOI might extend the infection dynamics and help to reveal the host response. Abundances of algal cells and extracellular viral particles were monitored for 23 days through the infection and recovery phases by flow cytometry (Supporting text). After 23 days, the recovered cultures were diluted weekly with fresh medium. The second infection assay included two exponential phase RCC6946 cultures; one was infected with EhVM1, which resulted in the recovery of a resistant culture. Growth curves were generated with GraphPad Prism 10.3.1 (GraphPad).

### RNA extraction and quality evaluation

Pellets for RNA extraction were collected by 2-step centrifugation at 4°C (Supporting text). Since the infection time series was done in the same flasks, we were limited in the available volume for RNA extraction. Depending on cell abundance, we pelleted 50-250 mL of each culture, where larger culture volumes (150-200 mL) were used when cell abundance was low. Cell pellets from uninfected treatment were collected until day 7, after the cultures reached stationary growth. Total RNA was extracted using the RNeasy Plant Mini Kit (Qiagen) according to the manufacturer’s protocol, followed by a double Turbo DNase treatment (Thermo Fisher Scientific, Invitrogen). RNA concentration and integrity were assessed using Qubit 2.0 Fluorometer (Thermo Fisher Scientific) and TapeStation assay (Agilent Genomics). During the advanced stages of infection (days 4-15), where live cell counts were low (10^3^-10^4^ cells mL^-1^), high-quality RNA extraction and library construction were unsuccessful (Fig. 1b bottom, empty circles). In the second infection assay, cell pellets for RNA extraction from a single time point during exponential growth were collected from *E. huxleyi* RCC6946 culture and a resistant recovered culture.

### MARS-seq library preparation and sequencing

cDNA libraries were prepared according to the bulk MARS-seq protocol (Keren-Shaul *et al*., 2019). A starting material of 50 ng clean RNA from each sample was reverse transcribed to cDNA using oligo(dT) adaptors containing molecular barcodes, allowing enrichment for mRNA and uniquely tagging each fragment. The tagged cDNA was pooled together, converted into double-stranded DNA (dsDNA), and linearly amplified using T7 RNA polymerase in vitro transcription (IVT). The amplified RNA was fragmented to 200-500 bp, ligated to partial Illumina primers, and reverse transcribed to cDNA before final amplification using 10-cycle PCR. The library size distribution and concentration were measured using Qubit and TapeStation assay. Quality control scores during library preparation were evaluated by qPCR of *actin* gene using 3’ directed primers: forward 5’-GACCGACTGGATGGTCAAG-3’; and reverse 5’-GCCAGCCTTCTCCTTGATGTC-3’ (Nam *et al*., 2018). cDNA libraries were sequenced using an Illumina NextSeq500 (75 cycles) or NovaSeq 6000 SP (100 cycles) platforms, aiming for 10M reads per sample. BCL files were demultiplexed based on molecular barcodes and converted to FASTQ files using the NGS pipeline UTAP (Kohen *et al*., 2019).

### Read processing and mapping to reference

Reads were assessed with FastQC (Andrews, 2010) and MultiQC (Ewels *et al*., 2016), trimmed with Cutadapt (Martin, 2011) to remove Illumina adapters and polyA tails, and filtered by Q20 quality and a minimal length of 25bp. The processed FASTQ files were downsampled to a depth of 10M reads and mapped to a dual reference of *E. huxleyi* and EhVM1 using STARv2.7.10a (Dobin *et al*., 2013) by the NGS pipeline UTAP (Kohen *et al*., 2019). The reference EhVM1 genome has 489 predicted CDS (Fromm *et al*., 2022), to which we added 306 potential ORFs (in total, 795 transcripts). Since *E. huxleyi* RCC6946 has no available genome, we used a pan-transcriptome built from several *E. huxleyi* strains (Feldmesser *et al*., 2021). The pan-transcriptome comprises 65,391 sequences and represents a broad repertoire of genes, comparable to genomes available for other *E. huxleyi* strains (Read *et al*., 2013; Skeffington *et al*., 2023; Kao *et al*., 2024). Gene annotations were obtained from Feldmesser *et al*., 2021, and additional KEGG annotations at the protein level from blastkoala (v3.1, eukaryotes database) and ghostkoala (v3.1, Eukaryotes + Prokaryotes + Viruses databases, score ≥ 100) (Kanehisa *et al*., 2016). Conserved domains were predicted using HHsuite version 3.3.0 (Söding, 2005) against the Pfam database (E-value < 0.05).

Genes encoding carbohydrate-active enzymes were predicted with the Carbohydrate Active Enzymes database (Drula *et al*., 2022) (CAZy, http://www.cazy.org/) based on *E. huxleyi* CCMP1516 draft genome (JGI Portal: Emihu1, NCBI Taxonomy ID: 280463). Annotated CAZy proteins of the draft genome (∼800) were blasted against the pan-transcriptome reference to convert JGI accession numbers to protein ID (BlastP, identity ≥ 95%, length ≥ 100). Cellular localization was predicted with Deeploc-2.1(Ødum *et al*., 2024) (default parameters for high-throughput analysis: high-quality model (slow) and short output format) with the recommended probability thresholds: cytoplasm (0.4761), nucleus (0.5014), extracellular (0.6173), cell membrane (0.5646), mitochondrion (0.6220), plastid (0.6395), endoplasmic reticulum (0.6090), lysosome/vacuole (0.5848), golgi apparatus (0.6494) and peroxisome (0.7364), Peripheral Membrane (0.60), Transmembrane (0.51), Lipid anchor (0.82), Soluble (0.50). Gene expression levels were evaluated with HTSeq-count v2.0.2 (Anders *et al*., 2015), and duplicate reads were removed using the Unique Molecular Identifiers (UMIs, molecular tags).

### Differential gene expression analysis

*E. huxleyi* genes with expression levels of ≥ 10 UMIs in ≥ 3 samples were considered significantly expressed and were included in the DE analysis conducted with the DESeq2 package (v1.38.3) (Love *et al*., 2014). Since RNA-seq provides compositional data representing relative abundances of genes, we excluded viral reads to prevent false identification of DE genes by introducing another organism (virus) into a finite pool of reads. Each time point under both conditions (infected and uninfected) was compared to the reference level of the uninfected sample at T0 (0.5 hours post-infection). Genes identified as significantly DE had baseMean expression ≥ 5, absolute log2(FC) ≥ 1 (Wald test: LFC threshold=1, alpha = 0.05, altHypothesis = “greaterAbs”), and an FDR adjusted p-value < 0.05 in at least one comparison.

For principal component analysis (PCA) and K-means clustering, we used normalized counts after variance stabilizing transformation (VST) using DESeq2 and R (v4.2.2). The optimal number of clusters was determined by complementary statistical methods. For heatmap visualization, we applied row (gene) standardization, scaling the means of each row to zero with a standard deviation of one.

The proportion of viral RNA transcripts was calculated out of the total transcripts mapped to the dual reference of *E. huxleyi* and EhVM1. Statistical significance was determined by a one-way repeated measures ANOVA and Dunnett’s post-hoc multiple comparison tests using GraphPad Prism 10.3.1 (GraphPad). Asterisks represent a statistically significant increase (p-value < 0.05). Some viral RNA transcripts exist in uninfected and recovered cultures (Fig. 2b). Since uninfected cultures were virus-free, the signal likely resulted from pipetting carry-over or incorrect assignment of reads to samples rather than viral expression. In the recovered cultures, the viral transcripts may reflect the background noise as uninfected cultures, suggesting recovered cultures are free of viral expression. Alternatively, they could originate from a small subpopulation that remains infected and expresses viral genes. This issue remains unclear, and future single-cell approaches could provide insights into it.

### Functional annotation and enrichment analysis

*E. huxleyi* Gene Ontology (GO) and KEGG annotations were adapted from Feldmesser *et al*., 2021. GO enrichment was computed by topGO package (v2.50.0) with Fisher exact test, weight algorithm, and biological process ontology. KEGG pathway enrichment was computed by applying a hypergeometric test in R. Enrichment of specific GO terms or KEGG pathways for each cluster was determined by p-value ≤ 0.01 when compared to all expressed genes.

### Meta-analysis of resistant strains

Published RNA-seq datasets comparing resistant and sensitive *E. huxleyi* strains were downloaded from Feldmesser *et al*., 2021, Kendrick *et al*., 2014, and Bousquet *et al*., 2025. The transcriptomes included four resistant *E. huxleyi* strains: CCMP373, CCMP379, LC5-11A (1N-flagellated), and LC5-12A (2N-flagellated). For CCMP373 (Feldmesser *et al*., 2021) and CCMP379 (Kendrick *et al*., 2014), processed FASTQ files were mapped to the dual reference of *E. huxleyi* and EhVM1 using RSEM (Li & Dewey, 2011). DE analysis was conducted with the DESeq2 package (v1.38.3). Data from Kendrick *et al*. had no replicates, so infected and non-infected samples were pooled together. For LC5-11A and LC5-12A (Bousquet *et al*., 2025), we used the available DE tables. Gene sequences from Bousquet *et al*. were compared to our pan-transcriptome reference to convert gene annotations (BlastN, identity ≥ 95%, length ≥ 100). In some cases, several genes in this database matched the same gene in the pan-transcriptome. Genes from each dataset were defined as DE when absolute log2(FC) > 1 and FDR-adjusted p-value < 0.05. From each dataset, DE genes that were shared with the time series transcriptome were organized according to the four clusters. Each gene was assigned a “conservation score” ranging from 1 to 5, reflecting the number of resistant strains in which it was upregulated. Similarly, downregulated genes were scored -1 to -5 based on the number of transcriptomes.

### Structural predictions and structural search

AlphaFold 3 (Abramson *et al*., 2024) was used to model the structure of the core resistant group genes and TIR-containing genes. Structural models were searched in the AlphaFold/UniProt50 database using the Foldseek webserver (van Kempen *et al*., 2024). Protein descriptions for each UniProt50 match were fetched using UniProt API. Top UniProt50 hits for each structural model were examined manually and further analyzed for functional annotation.

### TIR gene definition, phylogenetic, and structural analyses

To find TIR domain-containing proteins, conserved domains were predicted using HHsuite version 3.3.0 (Söding, 2005). TIR domain-containing proteins were defined as such if they contain at least one of the TIR-domain protein families: pfam13676, pfam08937, pfam01582, pfam10137, pfam08357, and pfam18567 (pfam probability score ≥ 50%). Expressed TIR domain-containing genes were manually examined for domain completeness and then structurally modeled as described above. Transmembrane helices were predicted using DeepTMHMM (Hallgren *et al*., 2022), and additional encoded domains were searched with the CDD database (Wang *et al*., 2023). PyMOL (v3.7; Schrödinger, LLC) was used to visualize the predicted structures and superimpose them with the human and Arabidopsis TIR domains.

## Supporting information

This PDF file includes:

Supporting text - extended Methods

Supporting Figures S1 – S6

Supporting Table S1

Supporting Data File 1:

- Supporting Data S1 – *E. huxleyi* infection with EhVM1 time series transcriptome count table (raw and normalized counts).

Supporting Data File 2:

- Supporting Data S2 – significantly differentially expressed genes of the time series transcriptome, including gene annotations.
- Supporting Data S3 – Significant results of functional enrichment tests (GO terms and KEGG)
- Supporting Data S4 – additional information to table 1 of up-regulated genes in resistant *E. huxleyi* populations
- Supporting Data S5 – Comparative transcriptomics analysis (meta-analysis) to other resistant strains.
- Supporting Data S6 – Structural homologs based on Alphafold predictions for resistance-related genes.
- Supporting Data S7 – A summary table of TIR-domain containing genes (as presented in Fig. 4)

## Extended Methods

### Cell enumeration

Samples were analyzed using the CytoFLEX S Flow Cytometer (Beckman Coulter) to quantify algal cell abundance. Cells were quantified by plotting the chlorophyll autofluorescence (excitation: 488 nm, emission: 663 - 737 nm) versus forward scattered light (a measure for cell size). Gates were defined according to the properties of cells during exponential growth, with chlorophyll fluorescence intensity higher than 10^4^ arbitrary units (a. u.) (Fig. S6a,d; “Live Cells” gate). To quantify cell death, samples were stained with Sytox Green (Invitrogen), a nucleic acid stain that penetrates only permeable membranes (a property of dead or dying cells). Samples were stained at a final concentration of 1 μM in the dark for 30 minutes at room temperature and analyzed by flow cytometer (excitation: 488 nm, emission: 500 - 550 nm). An unstained sample was used as a control to eliminate the background signal. The proportion of Sytox-positive cells was determined from the total detected cells in the “Live Cells” gate (Fig. S6b,e). At least 20,000 cells were analyzed per sample for all flow cytometry analyses. In samples from advanced stages of the infection, when the population was demised almost entirely, only ∼1,000 cells were analyzed per sample. Data analysis was performed with the CytExpert program (version 2.3.0.84).

### Assessment of extracellular viral particles and infectious viral particles

To enumerate extracellular viral particles, samples were fixed with a final concentration of 0.5% glutaraldehyde for 30 minutes at 4°C, flash frozen in liquid nitrogen, and stored at -20°C until processing. Fixed samples were thawed and stained at a 5:195 (μL) ratio with the nucleic acid stain SYBR Gold (Invitrogen) diluted according to the manufacturer’s protocol (5 μL SYBR Gold in 50 mL of 0.22 μM filtered TRIS-EDTA buffer), then incubated for 20 min at 80°C. Samples were analyzed by the CytoFLEX S flow cytometer (excitation: 488 nm, emission: 500 - 550 nm) with at least 1,000 events analyzed per sample (up to 350,000 events in infected samples) (Fig. S6c,f).

The infectivity of EhVM1 was evaluated using the Most Probable Number (MPN) assay (Jarvis *et al*., 2010). EhVM1 was serially diluted to the point of absence (10^-7^) and inoculated to an exponentially growing *E. huxleyi* RCC6946 culture (placed in a 96-well plate). Growth of the infected *E. huxleyi* RCC6946 cultures was monitored daily for a week with a plate reader (Tecan Infinite 200 PRO) by measuring chlorophyll intensity (excitation 480 nm, emission 660 nm). The concentration of infectious viral particles was estimated by the number of positive wells (infected, lysed culture) and statistical probability tables.

### RNA extraction and quality evaluation

Cell pellets for RNA extraction were collected 3 h after the beginning of the light cycle by 2-step centrifugation at 4°C. Samples were centrifuged at 24,500g for 12 min (Thermo Scientific Sorvall LYNX Superspeed Centrifuge), and the supernatant was removed. Then, cells were resuspended in a small volume (∼1 mL) and centrifuged at 16,000g for 1 min (Eppendorf 5417R Centrifuge). Cell pellets were resuspended in 450 μL RLT lysis buffer (Qiagen) containing 1% β-Mercaptoethanol and stored at -80°C until RNA extraction. RNA extraction was performed using the RNeasy Plant Mini Kit (Qiagen) according to the manufacturer’s protocol, followed by a double Turbo DNase treatment (Thermo Fisher Scientific, Invitrogen). RNA concentration and integrity were assessed using Qubit 2.0 Fluorometer (Thermo Fisher Scientific) and TapeStation assay (Agilent Genomics).

## Supplementary material

**Figure S1.**
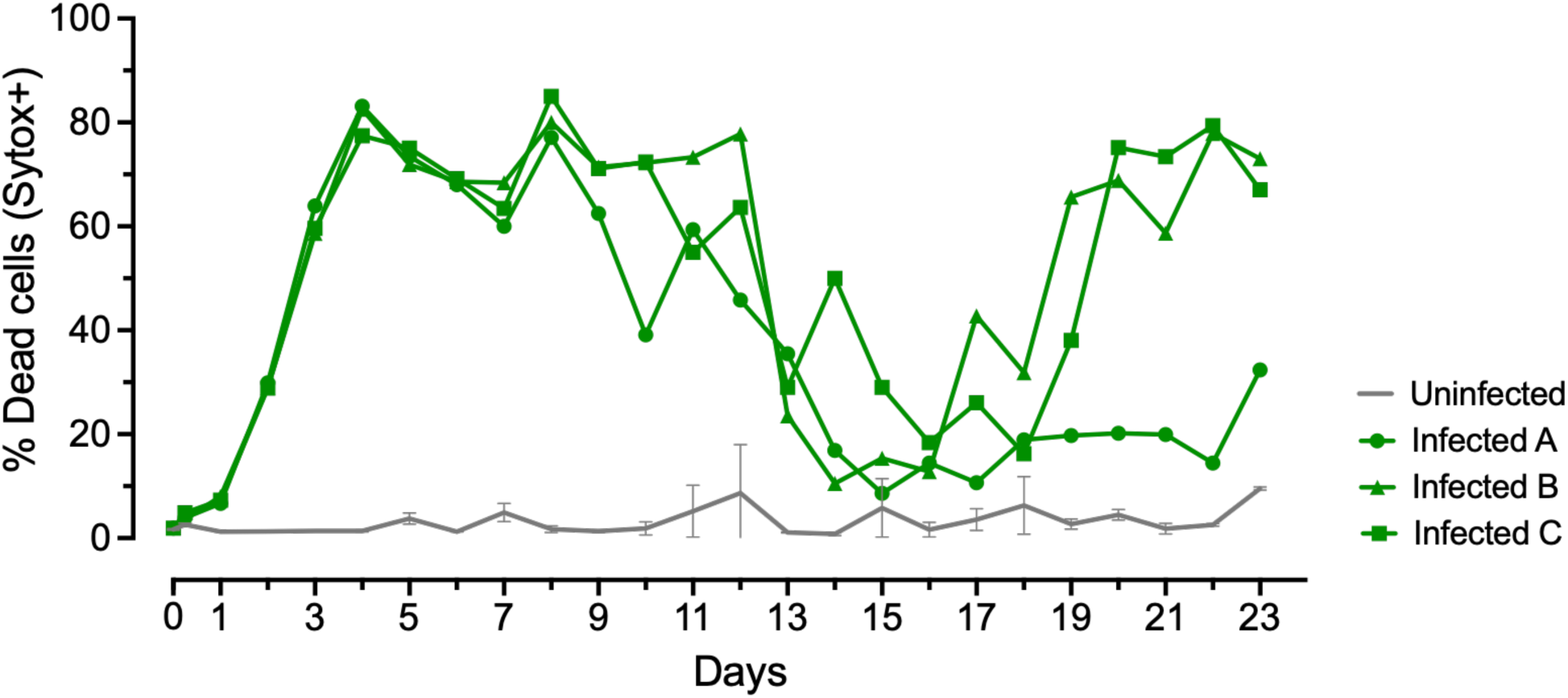
Cell death of *E. huxleyi* RCC6946 cells during EhVM1 infection. Cell death was measured by Sytox Green staining. Infected replicates are plotted individually to display the temporal variability. Uninfected replicates are represented as mean ± SD (*n* = 3). Error bars are clipped at the axis limit.

**Figure S2.**
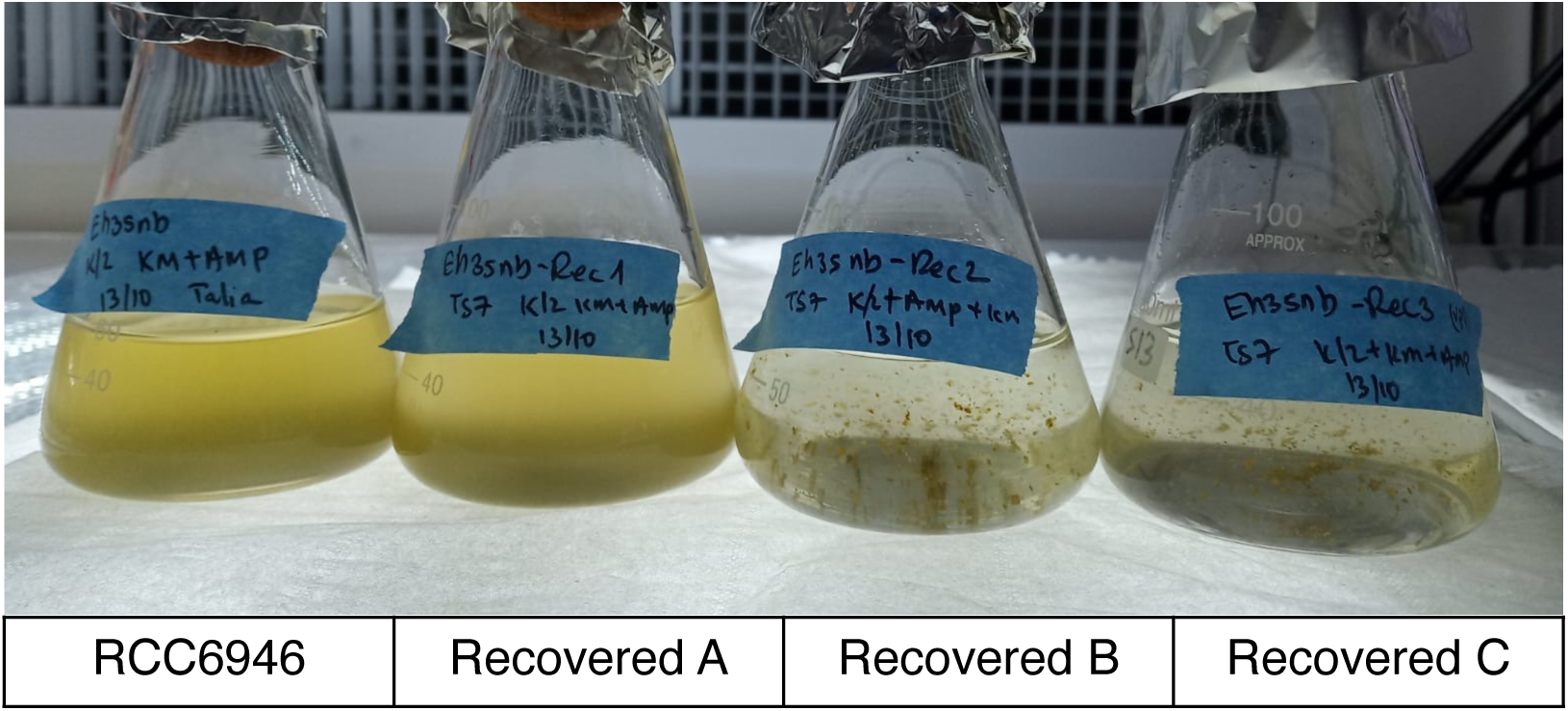
**Late recovery of resistant cultures and formation of cell aggregates**. The three cultures shown here are the recovered strains (Recovered A-C), resulting from Infected A-C cultures Out of the three recovered cultures, Recovered B and C exhibited longer recovery dynamics but both successfully recovered by day 37. After recovery, the two cultures formed green aggregates during exponential growth, while the Recovered A culture remained uniformly transparent green. The formation of aggregates is typically observed during viral infection and for cultures that coexist with the virus (Joffe *et al*., 2024).

**Figure S3.**
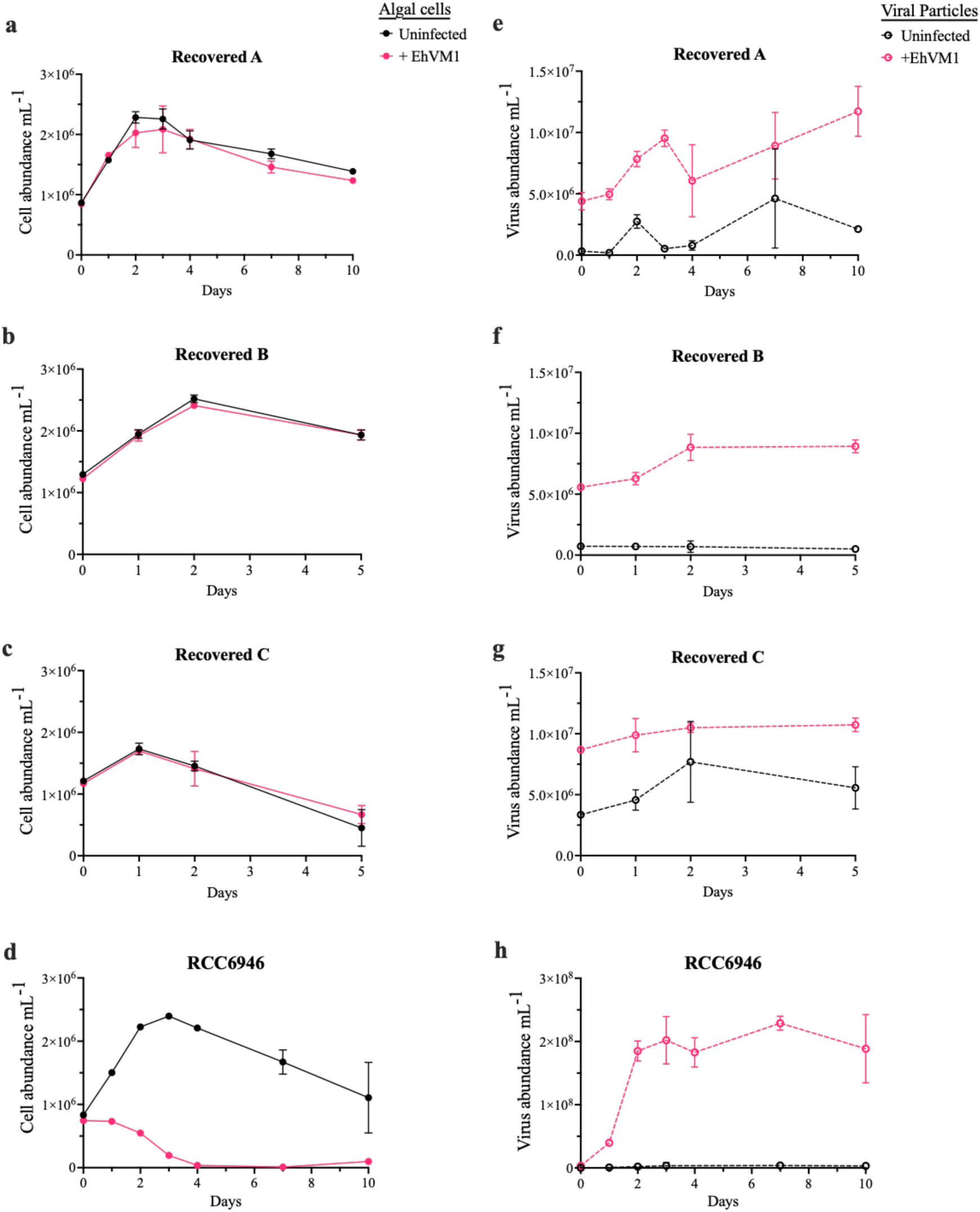
Infection dynamics of recovered strains and the ancestor *E. huxleyi* RCC6946 with EhVM1 to validate resistance. Cell and virus abundances were monitored by flow cytometry. Pink lines represent cultures to which we added viruses. (a-d) Cell abundance of three recovered populations and the ancestor *E. huxleyi* RCC6946. (e-h) Virus abundance. Note that the scale of EhV particles is different in (h) due to high viral production in infected *E. huxleyi* RCC6946. The detected viral particles in the recovered cultures A and C, where no viruses were added (shown by the dashed black line), are probably a result of carryover from the initial time series infection. Values are mean ± SD (n = 3). Error bars may be smaller than the symbol size.

**Figure S4.**
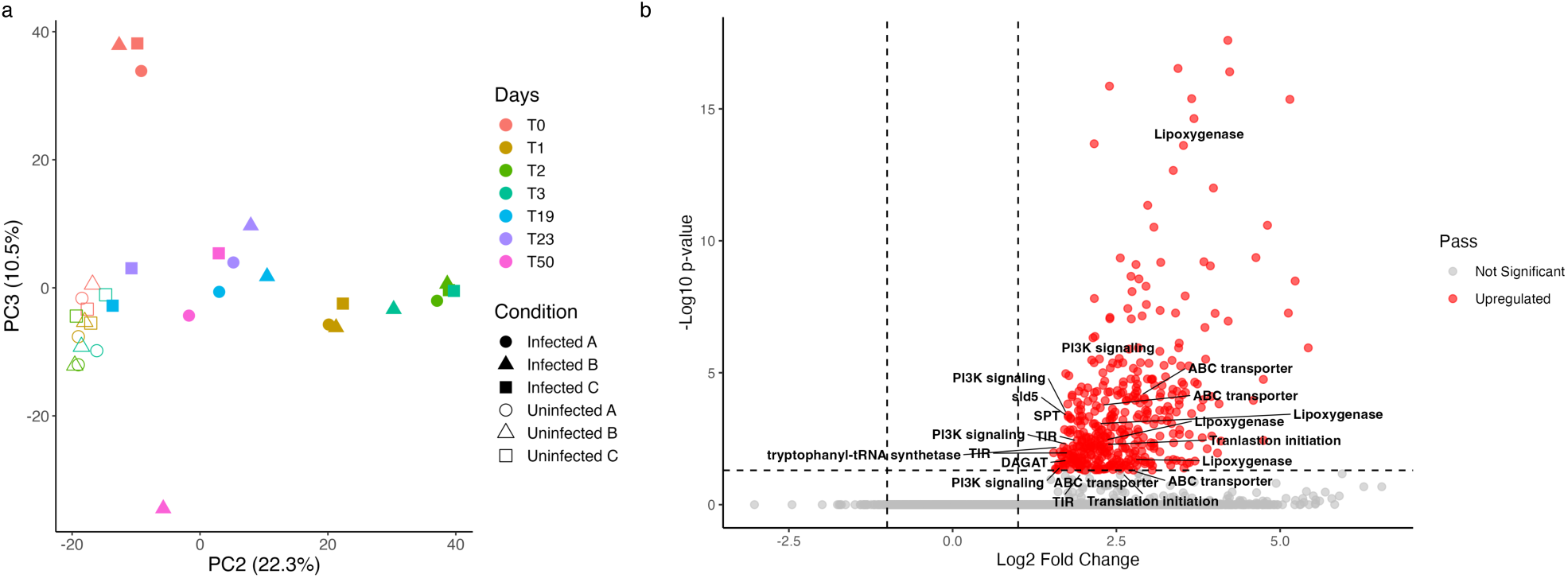
Early host response to infection. (a) PCA based on host gene expression values (normalized counts after variance stabilizing transformation, VST), focusing on PC3, which distinguishes the early response to infection at T0 (0.5h post-infection). (b) Volcano plot of host differentially expressed genes between uninfected and infected samples at T0. We detected 323 genes significantly up-regulated solely at T0, delineating an early and specific response to infection. Differentially expressed genes that passed the thresholds (log2(Fold Change) > 1 and FDR adjusted p-value < 0.05) are shown in red. Selected genes are marked with a text label. TIR: Toll/interleukin-1 receptor; PI3K: Phosphatidylinositol 3-kinase; DAGAT: Diacylglycerol acyltransferase; SPT: serine palmitoyltransferase.

**Figure S5.**
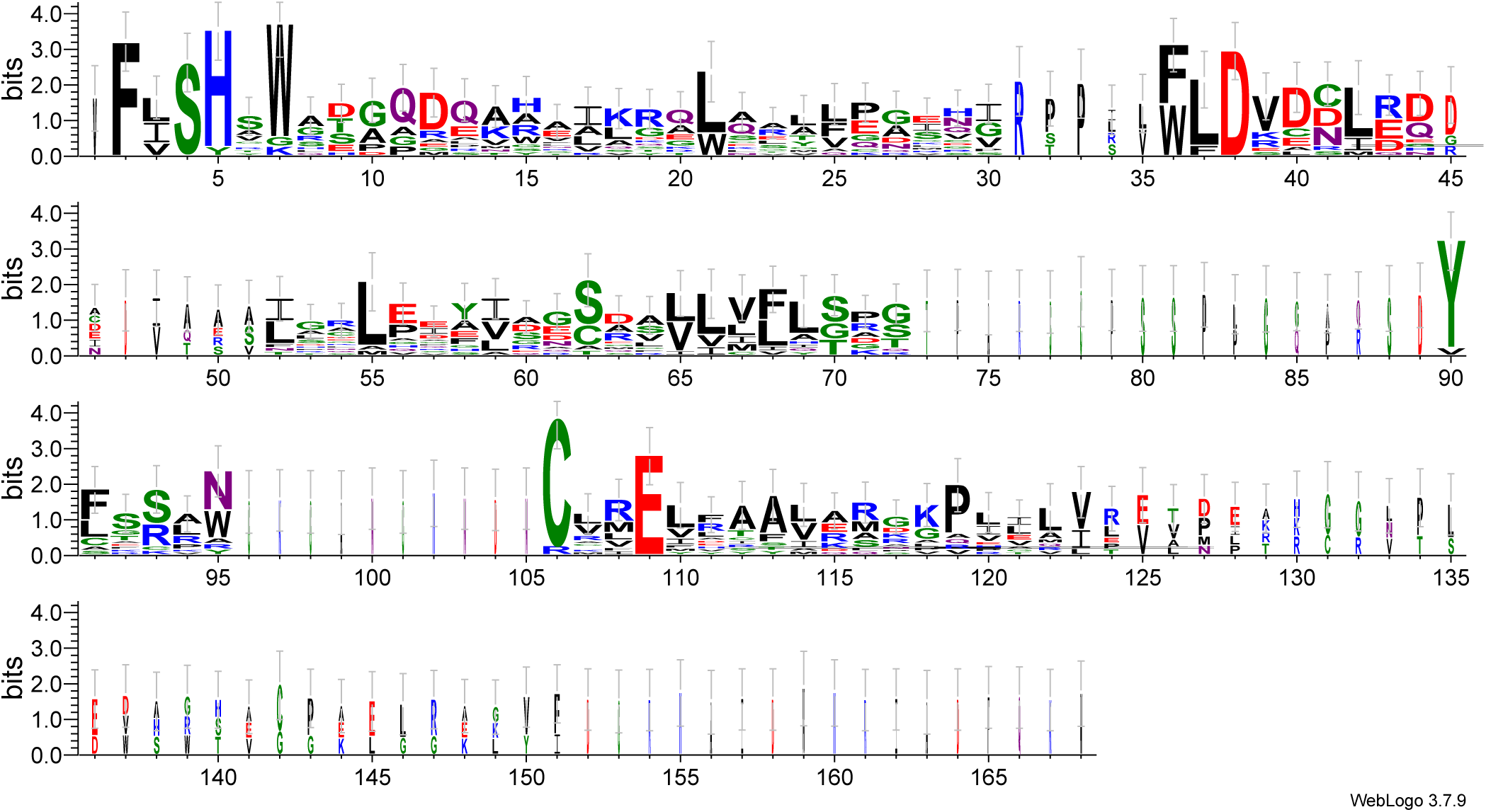
The putative catalytic glutamic acid residue is conserved in *E. huxleyi* TIR domains. The TIR-containing genes were searched in the CDD database to identify the location of the predicted TIR domain in each gene. Multiple sequence alignment on the domain part was performed using MUSCLE (v3.8.31) (Edgar, 2004) and visualized with WebLogo 3 (https://weblogo.threeplusone.com/) (Crooks *et al*., 2004) to detect conserved sequences or amino acids across the domain. The domains vary along most of the protein sequence, but several positions are conserved, specifically the aspartic acid D38 and the glutamic acid E109.

**Figure S6.**
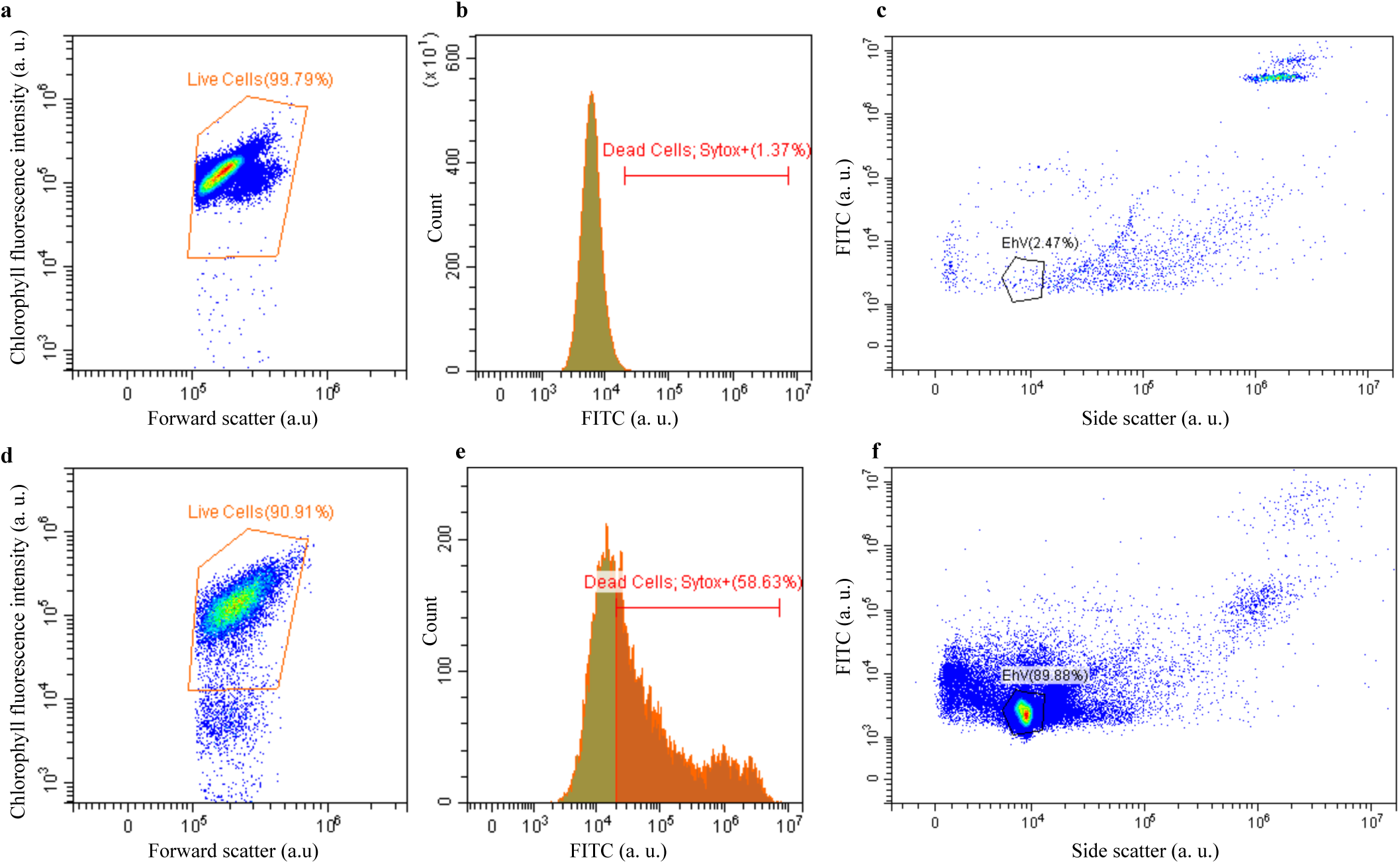
Quantification of *E. huxleyi* cell abundance, cell death, and viral particles. Flow cytometry measurements of (a-c) *E. huxleyi* RCC6946 cells during exponential growth (d-f) *E. huxleyi* RCC6946 cells infected with EhVM1 (3 days post-infection). (a, d) Dot plot analyses of cells based on their forward scatter area and chlorophyll fluorescence. (b, e) Histogram of Sytox+ stained cells, representing dead cells. Population gating was determined based on unstained control. Cells included in the Sytox analysis were derived from the “Live Cells” gate in a and d. (c, f) Dot plot representation of virus samples. The black gate marks EhV virions. Samples were fixed with glutaraldehyde and stained with SYBR Gold. a.u. = arbitrary units.

**Table S1.**
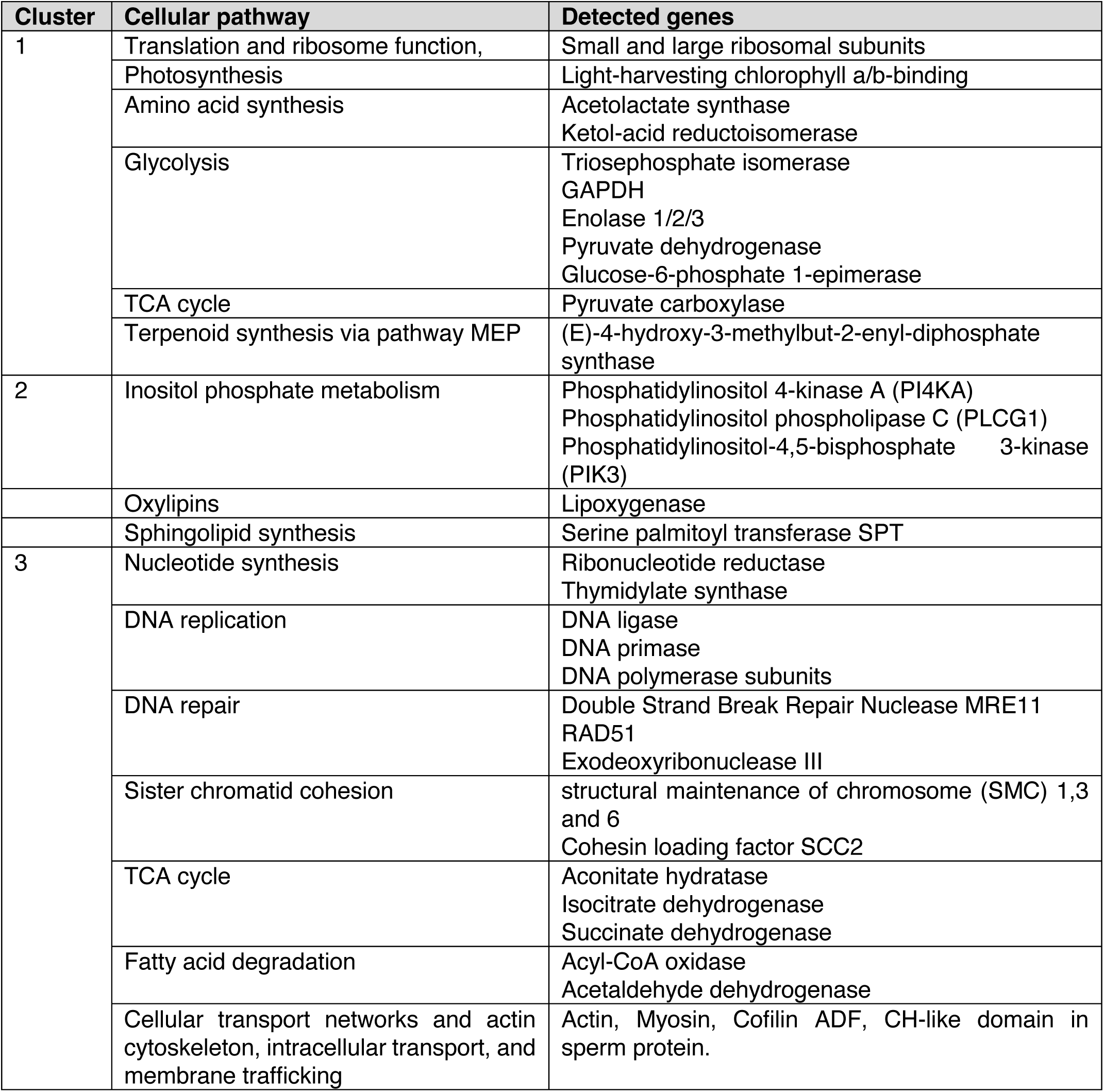
Overview of core metabolic pathways reprogrammed during infection of *E. huxleyi* populations.

